# Lipid Activation of a Thalamic GPCR Extends Working Memory Time-scales

**DOI:** 10.64898/2026.09.03.749183

**Authors:** James Newton Brandt, Alessandra Bonito-Oliva, Navid Paknejad, Celine Chen, John Fak, Andrew Luskin, Genelle Rankin, Miriam Kirylo, Leslie J Sibener, Amy F T Arnsten, Vanessa Ruta, Priya Rajasethupathy

## Abstract

Working memory capacity is poorly understood. The ability to control and extend working memory time-scales provides the potential to alleviate cognitive decline in disease and aging. Through an unbiased genetic search, we previously identified a thalamic orphan receptor Gpr12 as a potent enhancer of working memory, however its activation mechanism remains poorly understood. Here, we describe the CryoEM structure of Gpr12, revealing a lipidic regulatory site enabling an activated signaling state. By surveying the native lipidic environment in mouse thalamus we identified a class of 20 carbon:4 double bond fatty acid eicosanoids as potential ligands. Cell-based assays confirmed that the endogenous cannabinoid anandamide (AEA), but not other closely related family members or derivatives, robustly activates Gpr12. *In vivo* imaging during behavior revealed that Gpr12 activation produces a striking molecular state – the persistent suppression of cAMP in thalamus that tracks the duration of memory maintenance. Notably, genetic or pharmacological manipulations that enhance the AEA-Gpr12 signaling axis are sufficient to prolong cAMP suppression and extend the temporal window of memory maintenance. Furthermore, AEA-mediated cAMP suppressions in thalamus support sustained neural activity in PFC, specifically during memory maintenance. Thus, while cannabinoids often impair memory, here we identify an AEA-Gpr12 signaling axis in thalamus that enhances memory, including in primates. These findings identify a lipidic signaling mechanism in thalamus that is sufficient to control and extend working memory duration.

## INTRODUCTION

Working memory is a mental sketchpad used to temporarily maintain information long enough to perform goal-directed actions, such as reading the newspaper or planning dinner. While strongly linked to higher order executive function, it has long been appreciated that it is a limited capacity system^1^. The ability to maintain information in working memory declines over prolonged periods of time, spanning seconds to minutes, but it is not well understood what dictates the temporal limits of working memory maintenance. While classical studies have noted the involvement of the prefrontal cortex^2,3^ and other cortical association areas^4–7^ to support memory maintenance over seconds, emerging work suggest prominent contributions of the thalamus in sustaining cortical representations, particularly at prolonged periods of memory maintenance^8–12^. However, the molecular and cellular mechanisms that support these prolonged time-scales are unknown. Potential models have underscored the role of neuromodulation. For instance, the interaction between adrenergic, dopaminergic, or other GPCRs have long been predicted to generate persistent molecular signaling states^13,14^. But such molecular states that match the duration of memory maintenance, particularly at long time-scales, have not been identified. More importantly, mechanisms that can control or extend these time-scales are lacking. Thus, a deeper understanding of the mechanisms that set the temporal limits of working memory would not only be mechanistically powerful but would also be translationally meaningful, as working memory decline is prodromal and prognosis-driving in conditions such as ADHD, Schizophrenia, Alzheimer’s, and even natural aging.

Through a large-scale genetic mapping study, we previously identified a genetic locus in mice that accounts for substantial variation in working memory^11^. Within this locus, we identified an orphan receptor Gpr12, expressed exclusively in the brain, and functioning in thalamus, that can potently improve working memory maintenance. For instance, increasing thalamic Gpr12 expression through viral delivery was sufficient to improve memory retention across multiple tasks. Notably, Gpr12 is conserved in humans and is an emerging drug target for treatment of cognitive impairment^15,16^. These results motivate a search to identify the endogenous activating ligand of Gpr12, which can inform biological mechanism of memory duration, as well as rational drug design. Prior attempts to de-orphanize this receptor have resulted in conflicting reports, with proposed models ranging from various small molecule activators^17^ to self-activation of the receptor^18^. To date, no endogenous ligand has been identified.

To gain insights into the activation mechanism of Gpr12, we elucidated the cryo-EM structure of the receptor in its active state. This revealed a regulatory side pocket enabling the switch from a basal to potentiated state of the receptor. *In vivo* lipidomics followed by cell-based assays identified anandamide as a potent endogenous ligand driving this potentiated state. We found that thalamic synapses harbor the enzymatic machinery for on-demand anandamide synthesis and that activation of Gpr12 produces a prolonged molecular state-the persistent suppression of cAMP that scales with the duration of memory maintenance. Accordingly, anandamide delivery was sufficient to prolong cAMP depressions in thalamus, as well as neural activity persistence in PFC, specifically during periods of memory maintenance, thereby extending memory duration. Of note, these effects of anandamide were abolished in the absence of the receptor or with single point mutations that disrupt ligand-receptor interaction. Together, these findings identify an anandamide-Gpr12 signaling mechanism in thalamus that regulates the temporal window of working memory maintenance, and introduce the side ligand-binding pocket as a promising target for therapeutic development.

## RESULTS

### Gpr12 can achieve a ligand-gated activation state

GPR12 is among a small subset of GPCRs that exhibits constitutive activity in the absence of an exogenous ligand^19^. Such basal activity could arise from a ubiquitously present agonist occupying the orthosteric site or from intrinsic receptor features that stabilize the active conformation. A previous structure of GPR12 proposed a self-activation mechanism^18^ in which the extracellular loop 2 (ECL2) folds into the orthosteric pocket to act as a ligand, analogous to the mechanism described for GPR52^20^. However, our earlier observation that GPR12 has task-dependent functions during working memory^11^ led us to hypothesize the existence of a temporally gated activation mechanism not explained solely by receptor-intrinsic activity. We therefore sought to capture the active state of Gpr12 within a heterotrimeric complex consisting of Gαs, Gβ_1_, and Gγ_2_, by fusing its C-terminus to an engineered mini-G_s_^21^, which forces the receptor to remain in a ligand-binding conformation during expression and purification.

We obtained a 2.4 Å cryo-EM density map enabling unambiguous modeling of the complete GPR12 G-protein activation complex, excluding only the flexible 43-residue N-terminus and the synthetic Gly-Ser linker between GPR12 and miniG_s_ (Fig. 1A and 1D, Fig. S1-S3, Table S1). In this structure the orthosteric pocket of Gpr12 adopted an open conformation, with the pocket being occupied by a strong X shaped density, consistent with the amphiphilic detergent LMNG used during receptor purification (Fig. 1A-C). The acyl tails of LMNG extend deep into the orthosteric site, forming extensive hydrophobic contacts with a constellation of hydrophobic residues on TM1, TM3, and TM5 (Fig. 1B and 1C, Fig. S3B), while its sugar moieties are surrounded by polar and charged residues, with a substantial portion exposed to solvent. Notably, ECL2 does not occlude the orthosteric pocket, but instead cradles the LMNG molecule, separating its polar head groups and bulk solvent from the hydrophobic acyl tails extending into the pocket (Fig. 1B). This mode of ligand engagement is distinct from the ECL2-mediated self-activation architectures described for GPR52^22^, GPR21^23^, and GPR161^24^ (Fig. 1I).

**Fig. 1.**
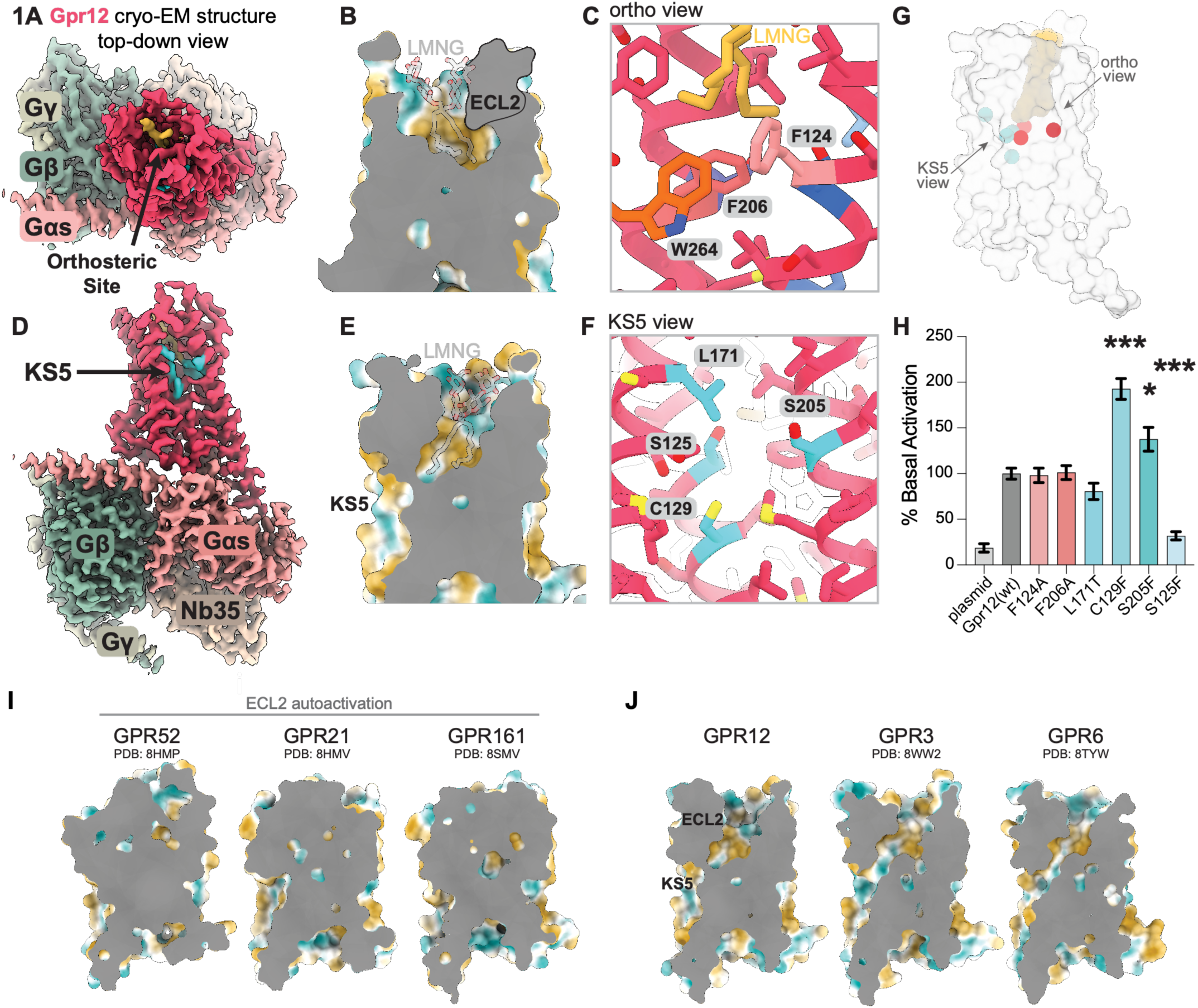
Gpr12 can achieve a ligand-gated activation state. (**A**) Top views of the 2.4 Å hGpr12-miniGαs-Gβ1-Gγ2 heterotrimeric signaling complex, stabilized by the nanobody Nb35. Gpr12 is shown in red, miniGαs in salmon, Gβ1 in green, Gγ2 in lime, Nb35 in light yellow, and orthosteric LMNG is shown in yellow (**B**) Cross-sectional view of orthosteric binding pocket with LMNG (transparent) relative to ECL2 (bolded black line) highlighting hydrophobic (yellow) and hydrophilic (blue) residues. (**C**) Orthosteric view inset, cross-section of the orthosteric pocket highlighting coordination by F124^3.36^ (salmon), F206 (red), and W264^6.48^ (orange) around LMNG (yellow). (**D**), Side views of the heterotrimeric signaling complex showing a second regulatory pocket (“KS5”) at the TM3-TM5 interface (blue). (**E**) Side cutout of orthosteric pocket with proximity to side pocket with hydrophilic residues in blue and hydrophobic in yellow with LMNG in transparent. (**F**) Side pocket view inset residues L171, S125, S205, and C129 (blues) shaping the pocket. (**G**) Distribution of point mutations (same coloring as c,f) relative to the orthosteric and side view. (**H**) *In vitro* receptor activity measured as cAMP accumulation in orthosteric-site mutants (reds), side-pocket mutants (blues) expressed as % over basal activity and normalized over Gpr12(wt). Plasmid denotes empty expression vector (pInducer21). Data are shown as mean +/− SEM (n=5-8), * p < 0.05, *** p< 0.001 One Way ANOVA with Dunnet’s test. (**I**) Visualization of the orthosteric pocket in receptors with a known self-ECL2 activation mechanism such as GPR52, GPR21, and GPR161. All three receptors have a continuous cryo-EM density throughout the orthosteric pocket. (**J**) Visualization of the orthosteric pocket in phylogenetically related GPCR relative side pocket with hydrophilic residues in blue and hydrophobic in yellow. In GPR3, GPR6 a channel between side and orthosteric pocket is continuous.

In addition to the orthosteric pocket, we observed that Gpr12 also contains a distinctive side pocket embedded in the membrane between TM3 and TM5 (Fig. 1D), sometimes referred to as KS5^25^ (Fig. 1E), exhibiting a cleft of hydrophilic residues (Fig. 1F). Such membrane-exposed side pockets of class A GPCRs have been described to bind fatty acids, and act as regulatory sites for Gpr40^26^ and recently for CB1R^27^. Notably, Gpr12 family members GPR3 and GPR6 share a continuous channel connecting the KS5 pocket to the orthosteric site (Fig. 1J), suggesting functional coordination, and a conserved structural feature that persists despite marked sequence divergence across Gpr12, Gpr6, and Gpr3 (Fig. S3C).

To assess how the orthosteric and KS5 sites contribute to receptor activity, we performed targeted mutagenesis of residues at both sites (Fig. 1G) and measured changes in Gpr12 signaling through cAMP production in a cell-based assay^28^ (Methods). As expected from previous reports^19,29,30^, wildtype cell surface expression of Gpr12 in HEK cells was sufficient to induce robust accumulation of cAMP (Fig. 1H; Fig. S4A-C). Interestingly, the introduction of single point mutations in the orthosteric site did not appreciably alter receptor activity. For instance, both Phe124 and Phe206 exhibit continuous density with LMNG, yet mutation of either residue to alanine did not alter cAMP levels (Fig. 1H). In contrast, mutations of residues within the side pocket led to large and bi-directional changes in receptor activity. For instance, substituting Ser125 with phenylalanine strongly attenuated receptor activity (Fig. 1H, without impacting cell-surface expression (Fig. S4B). More notably, substitution of either Cys129 or Ser205 to phenylalanine doubled receptor activation (Fig. 1H). Lastly, introduction of an Leu171T mutation, which alters local hydrophobicity, left receptor activity unchanged. Thus, mutagenesis of residues within the KS5 site identifies this cleft as a regulatory node that strongly and bi-directionally modulates Gpr12 activity. These findings suggest that rather than self-activation, Gpr12 is ligand-activated, and that the membrane embedded side pocket enables the switch from a basal to potentiated state of the receptor.

### Gpr12 is activated by the endogenous endocannabinoid anandamide

We next aimed to identify endogenous ligands that can activate Gpr12. Because Gpr12 functions in the thalamus during working memory, we hypothesized that analysis of the lipidomic milieu of Gpr12 in its native context would capture physiologically relevant lipid species. To map endogenous Gpr12 expression, we generated an epitope-tagged knock-in mouse (*Gpr12^3xHA^*) (Fig. S5A, S5B), enabling visualization of receptor expression and localization (Fig. 2A). To maximize the input material for lipidomic analysis, we also leveraged an epitope tagged viral over-expression construct of Gpr12, that when injected into thalamus matched the endogenous localization patterns (Fig. 2B), and remained functional, capable of improving working memory (reported previously in Ref 11). Using this construct, we performed large quantity immunoprecipitation of Gpr12, followed by lipidomic analysis.

**Fig. 2.**
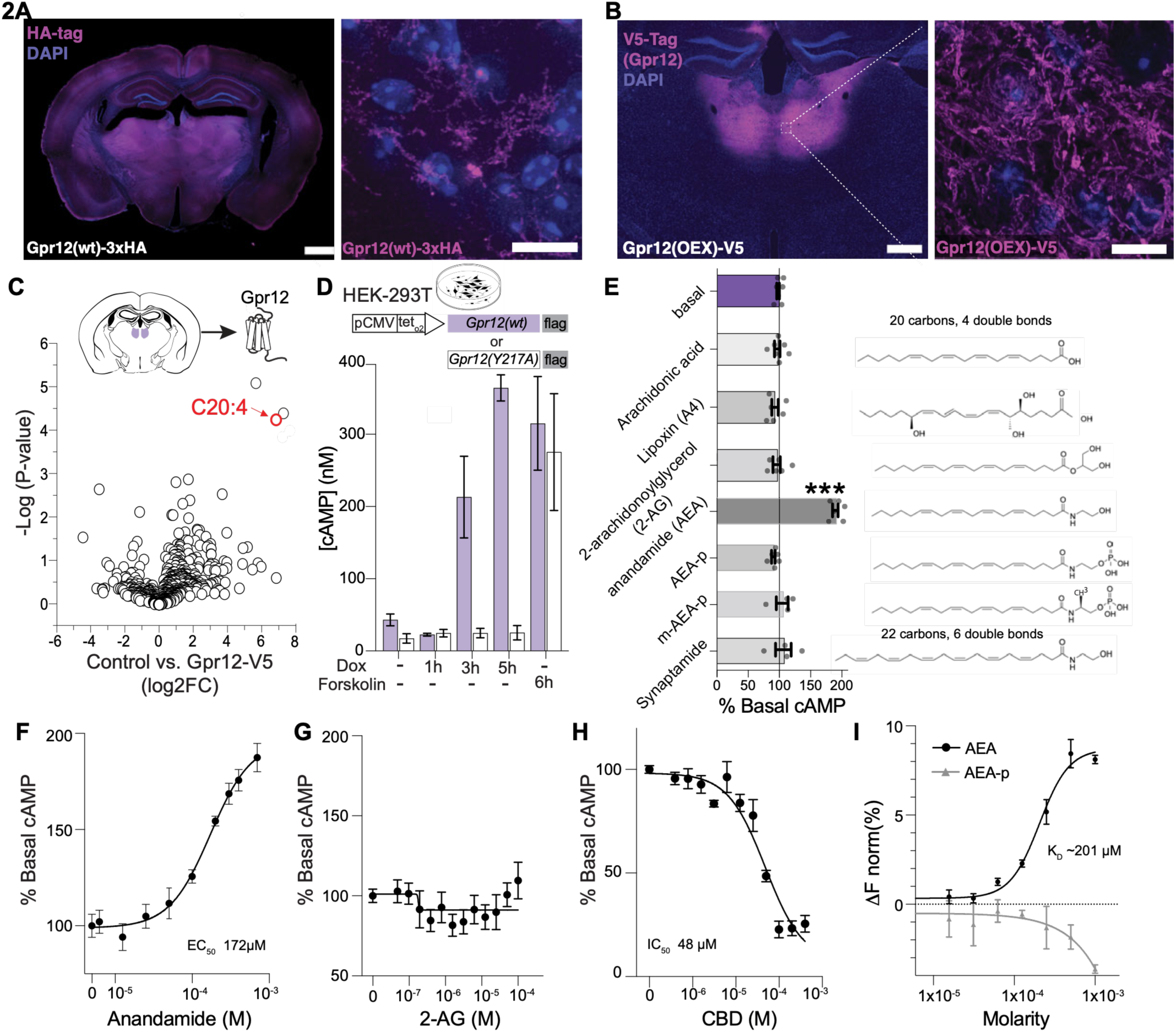
Gpr12 is activated by the endogenous endocannabinoid anandamide. (**A**) Coronal section from a Gpr12-3xHA mouse showing endogenous Gpr12 expression. Scale bars, 100 µm (left), 10 µm (right). (**B**) Thalamic pattern of viral Gpr12-V5 overexpression construct. Scale bars, 100 µm (left), 10 µm (right). (**C**) Unbiased negative-mode lipidomics (bottom panel) on Gpr12-v5 purified from the mediodorsal thalamus (top). 20 individual animals were pooled for each experimental condition (Gpr12-v5 and control). (**D**) *in vitro* cAMP accumulation with increasing doxycycline (Dox) incubation times (1hr, 3hr, and 5hr) in (Dox)-induced expression of wild-type hGpr12 (Gpr12(wt), purple) or constitutively inactive hGpr12(Y217A, white), or incubated with direct adenylyl cyclase activator forskolin. Data are mean +/− SEM from n=3 biological replicates. (**E**) Receptor activity measured as cAMP accumulation expressed as % over basal activity (0.1% DMSO). Arachidonic acid (n=7), lipoxin A_4_ (n=6), 2-arachidonylglycerol (2-AG, n=6), anandamide (AEA, n=6), anandamide-phosphate (AEA-p, n=6), methyl-anandamide-phosphate (m-AEA-p, n=4), and synaptamide (n=4), diluted at 300μM in 0.1% DMSO. Data are shown as mean +/− SEM (left) and correspondent chemical structures (right). *** p<0.0001, one-way ANOVA with Dunnett’s test. (F-H) Dose-response curve of Gpr12 activation induced by **F**) anandamide (3 replicates per condition), **G**) 2-AG (3 replicates per condition), and **H**) inhibition induced by cannabidiol (CBD) (3 replicates per condition). Data are shown as mean +/− SEM, normalized to basal activity (DMSO) condition. (**I**) MicroScale Thermophoresis binding assay showing change in fluorescent signal (670nm/650nm) in response to either anandamide (AEA) or anandamide phosphate (AEA-p). Data are 2 replicates per condition, shown as mean +/− SEM.

An initial unbiased analysis of the thalamic lipidome identified a distinct clade of lipids specifically enriched in Gpr12-v5 pull-down compared with control v5 pulldowns (Fig. 2C). Of the three mass-to-charge species within this clade, one exhibited a discernable fragmentation pattern consistent with an endogenous 20:4 eicosanoid (Fig. 2C, Fig. S5C). To assess the functional relevance of candidate 20:4 lipids, we developed a stable HEK cell line with Gpr12 induction under the control of a doxycycline-inducible promoter (Fig. 2D). We confirmed that increased levels of doxycycline-induced Gpr12(wt) led to progressive increases in cAMP levels, eventually paralleling forskolin-evoked levels as would be expected from its constitutive activity (Fig. 2D). As a control, we generated and expressed the DRY-motif mutant^31^ Gpr12(Y217A) to confirm that cAMP signaling depends on functional Gpr12. We further optimized doxycycline exposure and cell density to position basal Gpr12 signaling within the linear dynamic range of the assay^28^ (Fig. S5D), enabling detection of both positive and negative modulators. The assay was validated by recapitulating responses to a panel of previously reported exogenous inhibitors of Gpr12^19,32,33^ (Fig. S5E-H).

Using this assay, we performed a hypothesis-guided screen of neuronal eicosanoids^34–36^ and identified the 20:4 endocannabinoid anandamide as a robust activator of Gpr12, increasing receptor activity to ∼200% above basal levels (Fig. 2E). Other closely related eicosanoids (arachidonic acid, lipoxin A_4_), cannabinoid family members (2-AG), or molecules bearing slight modifications of anandamide (AEA-phosphate), had no activity on Gpr12 (Fig. 2E). Indeed, a systematic variation of acyl-chain length and head-group chemistry revealed that Gpr12 selectively recognizes lipids bearing an ethanolamine headgroup and a 20:4 acyl chain for robust potentiation (Fig. 2E). Modifying either feature-for instance extending the chain length as in synaptamide (22:6) or altering the headgroup to a phosphate (anandamide phosphate) or glycerol (2-AG,) – abolished activity (Fig. 2E). Examination of the anandamide dose-dependence revealed an estimated affinity of 172 μM at EC50 (Fig. 2F), which while within the affinity range reported for other lipid sensing GPCRs, including the phylogenetically related Gpr3-oleic acid pairing^37^, may also be an underestimate reflecting the limited aqueous solubility and strong membrane partitioning characteristic of lipidic ligands. Notably, the other major endogenous cannabinoid in brain, 2AG, had no activity on Gpr12 over a range of doses (Fig. 2G), while the exogenous cannabinoid (CBD) exhibited robust inhibitory activity (Fig. 2H). To assess whether anandamide directly engages Gpr12, we performed microscale thermophoresis binding assays using purified recominant receptor (Methods). Consistent with the functional screen, anandamide phosphate exhibited no detectable binding to Gpr12, whereas anandamide bound directly to Gpr12 with an apparent dissociation constant in range with the EC50 measured in cell-based signaling assays (Fig 2I). Collectively, these findings identify anandamide as an endogenous ligand in thalamus that is a potent and selective activator of Gpr12.

### Gpr12 elicits persistent cAMP depressions in thalamus that match memory duration

We next asked whether we can monitor Gpr12-elicited intracellular signaling in thalamus during working memory. Given the established coupling of Gpr12 to cyclic AMP, we developed an *in vivo* imaging approach to measure real-time changes in cAMP in thalamus as mice performed a working memory task. We tested several cAMP sensors and selected GFlamp2 for its high *in vivo* sensitivity. In brief, we injected mice with AAV9-GFLAMP2^38^ into the MD thalamus, and implanted a fiber optic cannula at the same site to record real-time fluctuations in cAMP fluorescence using a high sensitivity sCMOS camera, which we then aligned to ongoing animal behavior at tens of millisecond precision (Fig. 3A, Methods). We compared cAMP levels in AAV-Gpr12-mcherry (OEX) vs AAV-mCherry (control) groups to identify neural correlates associated with Gpr12-signaling and working memory enhancement (Fig. 3A-D).

**Fig. 3.**
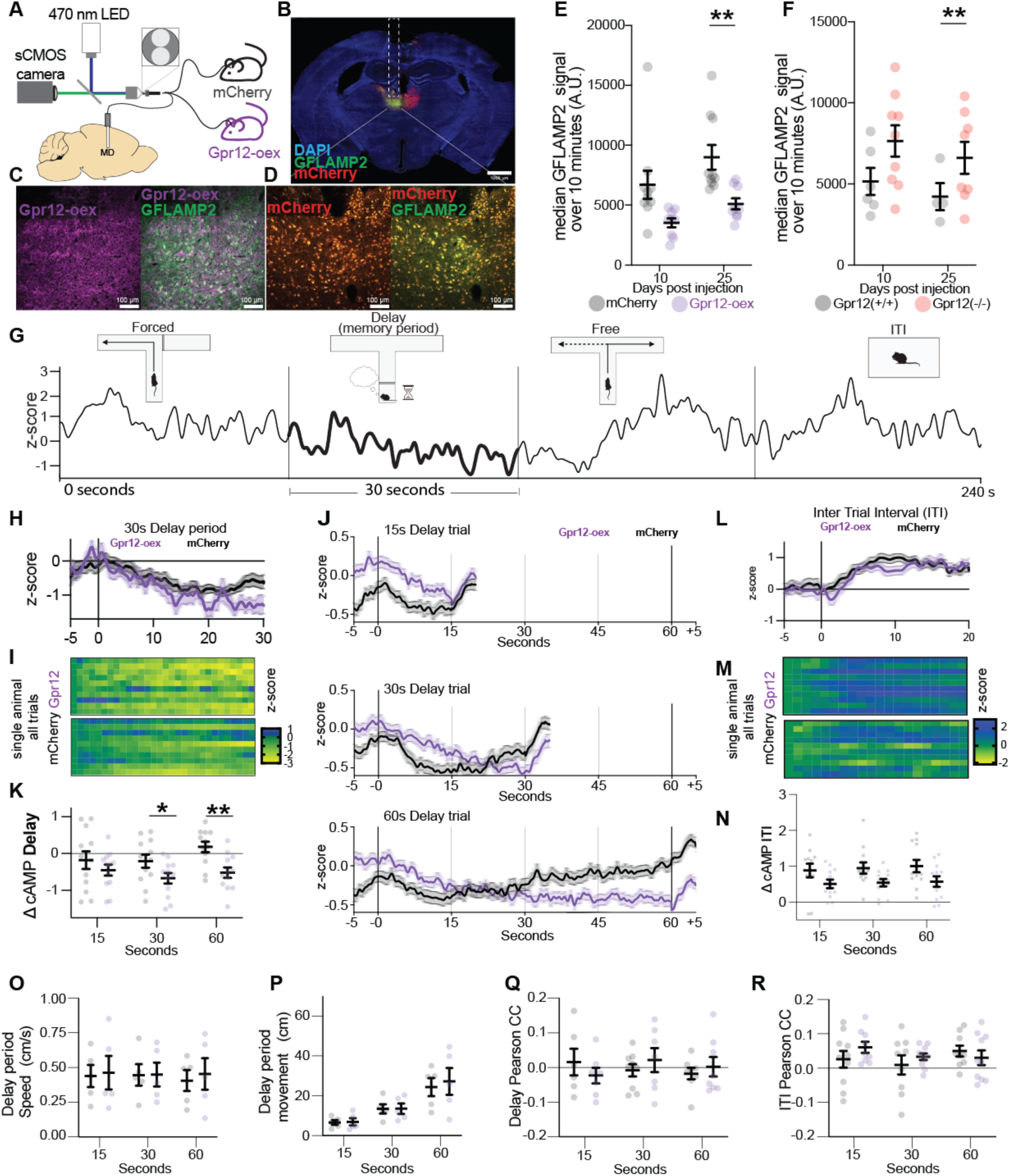
Thalamic Gpr12 drives persistent depressions of cAMP during memory periods. (**A**) Light path and schematic of the custom fiber-photometry system enabling simultaneous recording of cAMP signals through patch cords. Two mouse cohorts,Gpr12-oex (purple) and mCherry controls (black/grey), were recorded in parallel. (**B**) Coronal sections of viral expression in the mediodorsal thalamus. Bilateral mCherry expression with unilateral GFLAMP2 and cannula placement (white dash); Scale bars: 1000 µm (coronal) 4x stich, cannula diameter, 400 µm. (**C**) Insets show neuron-by neuron co-localization of Gpr12-oex expressing animals (purple) with GFLAMP2 in green. Scale bars: 100 µm, 20x confocal. (**D**) Insets show neuron-by neuron co-localization of mCherry expressing animals (red) with GFLAMP2 in green. Scale bars: 100 µm, 20x confocal. (**E**) Steady-state baseline cAMP recordings in mCherry control (n=10, grey) and Gpr12-oex (n=9, purple) animals. Data are mean +/− SEM, Welch’s T test, ** p< 0.01. (**F**) Steady-state baseline cAMP recordings in Gpr12 wild-type (Gpr12(+/+), n=6, grey) and Gpr12 knockout (Gpr12(−/−), n=8, orange) mice. Data are mean +/− SEM, Welch’s T test, ** p< 0.01. (**G**) Schematic of the delay non-match-to-place task, comprising the Forced, Delay (memory period), Free, and inter-trial interval (ITI) epochs. Below, representative single-trial cAMP traces from a Gpr12-oex mouse aligned to each task phase. (**H**) Trial-averaged, z-scored cAMP traces from a representative mCherry mouse (grey) and Gpr12-oex mouse (purple) across ten 30-s delay trials. Baseline differences in z-score values at start of delay period reflect differences in session means between groups (driven by delay period depressions in the oex group) rather than absolute differences in cAMP levels. Data are shown as mean +/− SEM. (**I**) Single-trial heatmaps from the same mice, with yellow indicating negative inflections (≤ –2 z). (**J**) Overlays of all trials from all mice at 15s (top), 30s (middle), and 60s (bottom) delay periods for Gpr12-oex (purple) and mCherry (grey). Data are mean +/− SEM, from mCherry and Gpr12-oex mice (n=12 each). (**K**) Change in cAMP during 15-, 30-, and 60-s delay periods, quantified as change in z-score from first 250ms to last 250ms of delay period. Data are mean +/− SEM from mCherry and Gpr12-oex mice (n=12 each), Welch’s test * p < 0.05, ** p< 0.01. (**L), (M), (N**) Analyses in D-F applied to the inter-trial interval (ITI) period. (**O**), (**P**) Quantification of distance moved (**O**) and velocity during delay periods (**P**) of DNMP task at 15, 30, and 60s delays. Data are mean +/− SEM from mCherry and Gpr12-oex animals (n=12 each). (**Q**) (**R**) Pearson’s correlation coefficient between the GFLAMP2 photometry signal and running speed during the delay period (**Q**) and the inter-trial interval (ITI) (**R**) in mCherry and Gpr12-oex animals (n=12 each). Data are mean +/− SEM.

In homecage baseline recordings, we found that Gpr12 OEX led to a significant reduction in steady-state cAMP levels in thalamus, as measured by median G-FLAMP2 fluorescence intensity compared to controls (Fig. 3E), raising the possibility that Gpr12 couples to Gα_i/o_ *in vivo*. However, these effects may also be due to underlying changes in sensor expression or receptor compensation. Since baseline fluorescence intensity measurements are subject to confounds related to sensor expression levels, we carefully titrated sensor expression, and used an isosbestic fluorescence measurement and post-hoc histological analysis to verify similar levels of sensor expression across groups, and that Gpr12 did not affect sensor expression (Fig. 3B-D, Fig. S6A). We additionally measured cAMP signals in Gpr12-knockout (−/−) animals, which exhibited higher baseline cAMP levels than WT (+/+) littermates (Fig. 3F, Fig. S6B), further indicating a basal inhibitory role for Gpr12 rather than compensatory effects. Thus, while we and others have observed Gpr12 coupling to Gs in HEK cells, that Gpr12 may couple instead to Gα_i/o_ *in vivo* in thalamus was functionally revealing, and consistent with other reports^39,40^ as well as our proteomic studies where Gpr12 favors proximity to Gi/o over Gs in thalamus (Fig. S9D), underscoring that GPCRs display distinct coupling and signaling depending on cellular context^41^.

To explore cAMP dynamics during a working memory task, we recorded cAMP levels as mice performed a widely used delay non-match-to-place task (DNMP), in which animals must remember a cue over gradually extending delays (memory periods lasting 15, 30, or 60s), to receive a food reward (Methods). We imaged cAMP signals in mice during all behavioral epochs, including the Cue, Delay (memory maintenance period), Choice, and inter-trial interval (ITI) periods (representative single trial trace of the cAMP signal shown in Fig. 3G). Across all trials (n=360), from both Gpr12-OEX and mCherry control animals (n=12 each), we detected no significant differences between groups in cAMP dynamics during the Cue, Choice, or ITI periods. However, specifically during the memory maintenance period, we identified a continuous cAMP depression, that was prolonged in Gpr12-OEX mice, matching the duration of memory maintenance (Fig. 3H representative mouse, averaging all trials; individual trials shown in Fig. 3I). More strikingly, when analyzing independent interleaved trials differing in memory duration (15, 30, or 60s), while control animals maintained a downward ramp for 15s, but not reliably at 30 or 60s, Gpr12OEX mice exhibited persistent downward ramps that scaled with the duration of memory maintenance (Fig. 3J, quantified in Fig. 3K), matching their improved behavioral performance at longer delays (described previously in Ref. 11 and also shown in Fig. 4H). Notably, these cAMP ramps returned to baseline immediately at delay-period offset (Fig. 3J), and the pseudo-random interleaving of trials ensures that the cAMP signals do not reflect anticipatory behavior but rather on-trial memory duration. Such differences in cAMP dynamics between OEX vs controls were not observed during the ITI period, the other largely stationary phase of the task of similar duration and animal motion (Fig. 3L-N). Furthermore, the observed delay-period differences in cAMP were not due to differences in locomotion during the delay period (Fig. 3O-R, Fig. S6C,D). These data identify that real-time cAMP depressions in thalamus serve as a well-matched neural correlate of working memory maintenance. They further support a model of basal Gpr12 activity in thalamus (chronic cAMP suppression at rest, Fig. 3E-F) juxtaposed with periods of activity-dependent modulation (task-related acute cAMP depressions, Fig. 3J) supporting context-dependent engagement during memory periods.

**Fig. 4.**
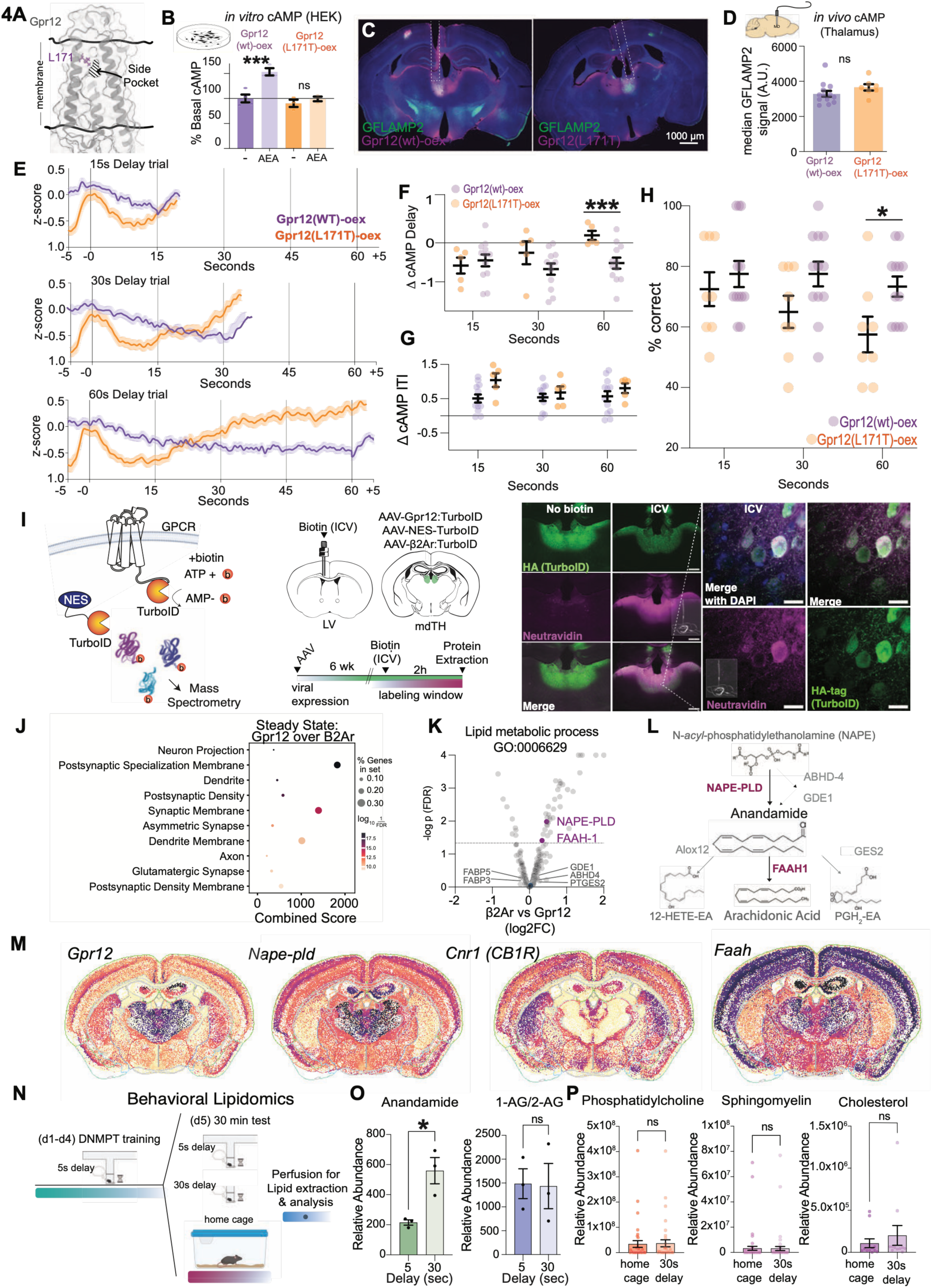
Disrupting AEA-Gpr12 signaling shortens cAMP depressions and memory duration. Membrane-facing side view of Gpr12 with L171 highlighted (purple) in close proximity to the side pocket. (**B**) HEK cell cAMP accumulation following 100 µM AEA challenge in Gpr12(wt)-oex and Gpr12(L171T)-oex; data are mean +/− SEM (n=3/each). (**C**) Coronal sections of viral expression in the mediodorsal thalamus showing Gpr12 expression with Gpr12(wt)-oex (left) and Gpr12(L171T) (right), purple, with unilateral GFLAMP2 (green) and cannula placement (white dash); Scale bars: 1000 µm (coronal) 4x stich, cannula diameter, 400 µm. (**D**) Steady-state photometry cAMP measurements comparing Gpr12(L171T)-oex, orange with Gpr12(wt)-oex, purple, 25 days after viral injection. Data are mean + SEM from Gpr12(wt)-oex (n=12, purple) and Gpr12(L171T)-oex (n=8, orange). (**E**) Photometry cAMP signal from Gpr12(wt)-oex and Gpr12(L171T)-oex during the delay non-match to place working memory task. Overlays of all trials at 15s (top), 30s (middle), and 60s (bottom) delay periods for Gpr12(wt)-oex (purple) and Gpr12(L171T)-oex (orange). Data are mean +/− SEM. (F-G) Change in photometry signal from Gpr12(wt)-oex and Gpr12(L171T)-oex from beginning to end of the delay period (**F**) and the ITI (**G**) of the delay non-match to place working memory task. Data are mean +/− SEM, Welch’s t test *** p< 0.001. (**H**) Short-term memory performance expressed as % Correct choices in a DNMP task. Data are mean +/− SEM from Gpr12(L171T)-oex (n=8, orange) and Gpr12(wt)-oex (n=12, purple), Welch’s t test *p< 0.05. (**I**) Cartoon of TurboID-mediated proximity labeling of thalamic GPCR-TurboID constructs (left) and schematic of experiment (center) showing protein extraction performed 6 weeks after viral injection and 2 h following ICV biotin delivery for thalamic proteome comparisons. Histology of Gpr12-TurboID expression and biotin-labeled proteins (streptavidin) in primary hippocampal neurons. Scale bar is 10 µm, 63x confocal (right). (**J**) GO-term enrichment of Gpr12 vs B2AR proteomes highlighting subcellular localization of Gpr12. (**K**) TurboID-mediated proximity labeling of thalamic GPCR-TurboID constructs from n=3 samples each of four pooled animals, volcano plots compare proximal proteomes of b2AR and Gpr12, highlighting GO:0006629 lipid metabolic proteins. (**L**) Metabolic (NAPE-PLD, ABHD-4, GDE1) and Catabolic (FAAH, Alox12b, PTGES2) anandamide generation enzymes with identified Gpr12-proximal proteins in purple. (**M**) MERFISH maps showing thalamic enrichment and overlapping spatial gene expression profiles of Napepld and Gpr12, Brain Knowledge Platform, Allen Institute. Signal intensity is represented as log2 (CPM+1) with purple indicating higher transcript abundance and beige indicating lower transcript abundance(**N**) Schematic of the behavioral lipidomics experiment. After 4 days of DNMP training, mice were randomly split in three groups and the following day were either subjected to a 30-minute test consisting of either 5 or 30 second delays or taken from the home cage, and successively perfused for lipid extraction and analysis. (**O**) Targeted lipidomics analysis of anandamide (green) and arachidonic acid (1-AG/2-AG) (blue) after a 5s (dark) and 30s (light) working-memory test. Data are shown as mean +/− SEM (n= 3 biological replicates). Welch’s test * p < 0.05. (**P**) Untargeted lipidomics analysis showing Phosphatidylcholine (left), Sphingomyelin (center) and Cholesterol (right) expressed as relative abundance in home cage and 30s delay group. Relative peak area denotes anandamide or arachidonic acid normalized to d31-palmitate (n= 3 biological replicates).

### Disrupting AEA-Gpr12 signaling shortens cAMP depressions and memory duration

We next sought point mutations that disrupt AEA-Gpr12 signaling to assess the causal effects on cAMP dynamics. Since the *in vivo* signaling features of chronic cAMP suppression at rest and acute depression during memory maintenance, closely parallel the basal and potentiated receptor states revealed by our structural analyses, we hypothesized that the functionally relevant transition between these states may be governed by anandamide activation of Gpr12 via the KS5 side pocket. We therefore searched for point mutations that selectively uncouple anandamide-evoked potentiation from basal receptor activity. We found that a single substitution in KS5, the L171T mutation (Fig. 4A; 1H), was sufficient to preserve constitutive cAMP signaling in HEK cells while disrupting anandamide-evoked activation (Fig. 4B).

We virally expressed Gpr12(L171T) in thalamus, comaprable in expression to the Gpr12-OEX (WT) (Fig. 4C, Fig. S7A-C), and found in homecage recordings that Gpr12-OEX(L171T) was still able to elicit baseline cAMP suppression characteristic of Gpr12-OEX (WT) (Fig. 4D), suggesting that the constitutive state remains intact. However, during DNMP behavior, Gpr12(L171T) mice displayed a marked failure to sustain cAMP depressions during the memory maintenance period, compared to Gpr12 (WT)(Fig. 4E). These differences in cAMP dynamics for the L171T mice were unique to the delay period as there were no observable differences during similarly timed ITI periods (Fig. 4F,G). Critically, Gpr12(L171T) animals were able to learn the DNMTP task at the same rate as Gpr12(wt)-oex animals (Fig. S7D), yet exhibited diminished performance at 30s and 60s delay periods (Fig. 4H). These results demonstrate that a single point mutation in the KS5 pocket disrupting the anandamide-activation of Gpr12 is sufficient to disrupt cAMP suppression and memory maintenance.

We next sought to understand the localization and timing of AEA-Gpr12 interactions *in vivo*, and whether they could signal on the spatial and temporal scales required for activation during working memory. The mechanisms by which lipid ligands reach and engage their receptors in the brain remain poorly understood, with hypotheses ranging from cell type specific “on demand” synthesis^42,43^ to multiple proposed routes for membrane^44,45^, synaptic^46^, or intracellular transport^47^. To determine which of these mechanisms supports endogenous anandamide-Gpr12 coupling, we used proximity labeling to map the membrane proteome of Gpr12⁺ thalamic neurons. In this approach, we fused the biotin ligase TurboID to Gpr12 (Fig. 4I), such that, in the presence of biotin, it generates reactive biotin-AMP and biotinylates nearby proteins within 10-30nm^48^. Thus, the expression of Gpr12-turboID in thalamus together with local biotin delivery is expected to result in tagging of the local interactome by biotin.

We generated a functional Gpr12-TurboID construct (Fig. S8A-C) as well as one for the beta-adrenergic receptor Adrb2-TurboID (β2AR), which is another membrane-localized GPCR with an established proximity-labeling dataset^49^, to allow for the identification of Gpr12-specific interactions. We determined that standard intraperitoneal (IP) delivery of biotin led to weak penetration in brain, and we thus evaluated additional routes, identifying that intracerebroventricular (ICV) injection produced robust, temporally restricted biotinylation detectable by immunohistochemistry (Fig. 4I, S8D), silver staining (Fig. S8E), and western blotting (Fig. S8F,G). We virally delivered either Gpr12-turboID or Adrb2-turboID into mediodorsal thalamus, along with ICV biotin, extracted thalamic tissue, and processed the samples for Mass Spectrometry (Methods, Fig. S9A). We confirmed that known interacting proteins were present within the identified B2AR proximal proteome (Fig. S9B) and that the interactome pattern of Gpr12 was consistent with its post-synaptic localization in the thalamus (Fig. 4J, Fig. S9C). Furthermore, when evaluating the G-protein abundances within the proximal Gpr12 proteome, we found that GNAO1 (Gi/o) was preferentially enriched, consistent with downward cAMP activity observed in the photometry experiments (Fig. S9D).

We focused our analysis on the potential lipidic pathways present within the local interactome of Gpr12. Interestingly, we found no Gpr12-specific enrichment of proposed anandamide-shuttling proteins, including the catalytically inactive FAAH-1 variant^47^, FABP7, or FBP5, nor did we detect components of the ABHD4/GDE1 biosynthetic pathway or the anandamide-degrading enzymes ALOX12B or PTGES2. Instead, Gpr12⁺ membranes showed specific enrichment of NAPE-PLD and FAAH-1 (Fig. 4K, Fig. S9E), the canonical enzymes responsible for anandamide synthesis and hydrolysis, respectively^50,51^ (Fig. 4L). Analysis of an open-source MERFISH dataset^52^ confirmed mediodorsal thalamic enrichment of NAPE-PLD that mirrored the spatial profile of Gpr12 expression with an absence of the CB1R in the thalamus (Fig. 4M). Among the proposed routes for anandamide signaling, these data strongly support local production and turnover of anandamide near Gpr12, which can occur within the seconds-to-minutes level timescales of behavior^50,53,54^.

Finally, to assess whether lipid profiles in the thalamus are dynamically changing during a working memory task, we performed lipidomics before and after 30-min of cognitive engagement, where mice performed repeated trials of DNMP with minimal working memory component (5s delay periods) or high demand working memory (30s delay periods) (Fig. 4N). A targeted analysis for AEA revealed markedly increased levels specifically after high demand working memory trials (Fig. 4O), whereas the levels of 1-and 2-AG endocannibnoids, as well as broad categories of membrane lipids, largely remained the same (Fig. 4P). Taken together, these findings identify local anandamide production and engagement of the Gpr12 side pocket as a mechanism to regulate the persistence of cAMP signaling and memory duration.

### Enhancing AEA-Gpr12 signaling prolongs cAMP depressions and improves working memory across species

To test whether enhancing the AEA-Gpr12 signaling axis is sufficient to extend cAMP depressions and memory duration, we tested the effects of direct anandamide delivery, in the presence and absence of Gpr12. We first assessed the effect of i.p. anandamide delivery on an innate spatial memory task, in which mice naturally prefer to explore new arms of a 5-arm maze, and we assessed the percentage of unique 3-, 4-, and 5-arm entries as a measure of working memory. This is an extension of the commonly used Y-maze spontaneous-alternation task^55^, modified into a five-arm radial maze^56^ with distinct visual and tactile cues at each arm, enhancing exploration. We found that i.p. anandamide improved working memory across multiple difficulty levels, including unique 3-, 4-, and 5-arms, each requiring progressively longer memory maintenance periods (Fig. S10A), which were not due to changes in gross motor activity (Fig. S10B). Importantly, the anandamide-dependent memory enhancements required an intact Gpr12, as such effects were absent in Gpr12 −/− mice (Fig. S10A).

To further assess the robustness of these effects, and define the temporal relationship between Gpr12 activation, cAMP depression, and working memory, we tested the effects of anandamide in the DNMP task (Fig. 5A), where we can more precisely control memory duration. Here, we found that i.p. anandamide led to a sizeable extension in the duration of cAMP suppression during the delay period of the task, specifically at extended delays (Fig. 5B-E; representative mouse shown in Fig. 5B as average of all trials, within individual trials shown in Fig. 5C; Quantified over all mice in Fig. 5D,E). This was associated with significant improvements in memory retention on similarly extended time scales (p = 0.004, paired t-test, n = 12) (Fig. 5G), which importantly was not present in Gpr12 −/− mice (Fig. 5H). Specifically, when we tested Gpr12 −/− mice in this task, they were able to learn the task similarly to their controls (Fig. 5F), but exhibited pronounced deficits performing near chance at longer 60s delays, which notably was not improved by i.p. anandamide (Fig. 5H p = 0.19, paired t-test).

**Fig. 5.**
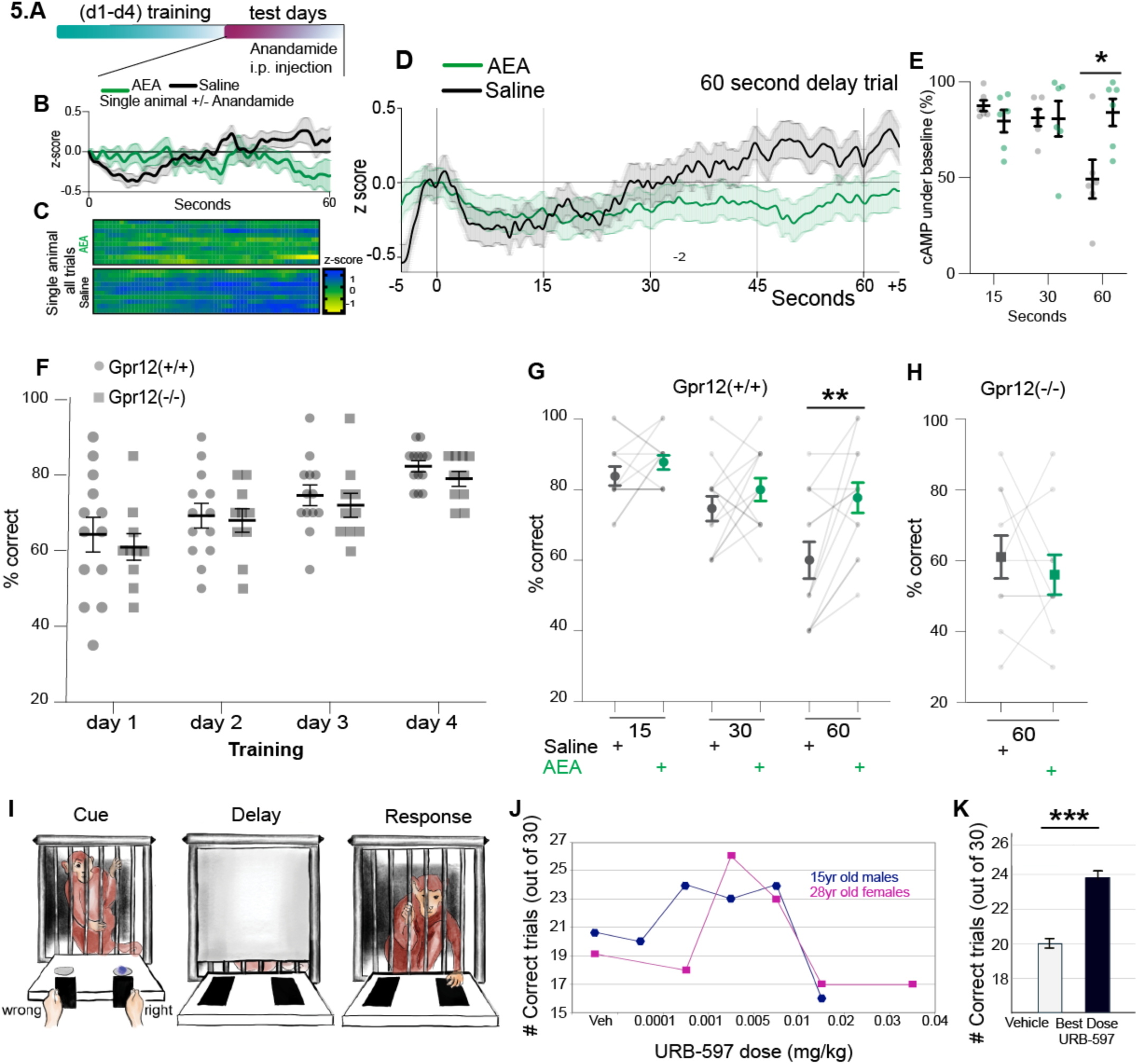
Enhancing AEA-Gpr12 signaling prolongs cAMP depressions and improves working memory across species. (**A**) Schematic of photometry recording in MD during the DNMPT test days following saline or AEA i.p. injection. **B**) Trial-averaged, z-scored cAMP traces from a representative mouse receiving saline (grey/black) or 5 mg/Kg anandamide (AEA, green) across ten 60-s delay trials on consecutive days. Data are mean +/− SEM. (**C**) Single-trial heatmaps from the same mice, with yellow indicating negative inflections (≤ –2 z). (Data are mean +/− SEM). (**D**) Photometry cAMP signal from Gpr12(wt) injected with saline or Anandamide during delay non-match to place working memory task. Overlays of all trials from all mice at 60 s delay periods for saline (grey) and Anandamide (green). (**E** Change in cAMP depression in mice receiving Anandamide (green) or saline (grey), quantified as the percentage of time the photometry signal remains below baseline (z=0). (**F**) Learning curve during the training phase of the DNMP task in Gpr12(+/+) (n=13, circle) and Gpr12(−/−) (n=10, square). Subsequently, each group is split in two and injected with either saline or 5 mg/Kg AEA to perform the test shown in Fig. 5 G/H. The test was repeated on the following day with the opposite treatment so that all mice were tested following saline and AEA injection. Two-way ANOVA with repeated measures, n.s. (**G**) DNMP test performance of Gpr12(+/+) mice (n=13) injected i.p. with saline (black) or AEA (green) during a 15, 30, and 60s delay test. Data are mean +/− SEM, paired t-test **p< 0.01. (**H**) Performance of Gpr12(−/−) mice (n=10) injected i.p. with saline (black) or AEA (green) during 60-s delay periods. Data are mean +/− SEM. (**I**) Task structure of the Visuospatial working memory performed with a manual response in a Wisconsin General Test Apparatus, consisting of a cued, a variably delay period “Delay” and a response period “Response” where the animal signals initial cue location. (**J**) Sample dose-response curves for intramuscular injection of the FAAH inhibitor, URB-597 administered intramuscularly 2 hrs before testing to two animals, a 15 year old male (blue) and a 28 year old female (pink) with Correct number of trials on Y axis and URB dose in mg/kg on X axis. (**K**) Grouped data from 14 total subjects of maximal effective URB-597 dose. *** p< 0.001, Paired T test.

Finally, to test for conserved effects of anandamide on working memory in primates, we delivered the FAAH-1 inhibitor URB-597 to rhesus macaques as they performed a visuospatial working memory task, which similar to the DNMP, consists of a cue, delay (0-60s), and choice period. We tested 14 macaques, and performed dose-response curves in each, to identify doses of maximal effect. Strikingly, we found that pharmacological elevation of endogenous anandamide, via delivery of the FAAH-1 inhibitor URB-597, robustly improved working memory in rhesus macaques, over time-scales of up to a minute (P<0.001, n=14 macaques, Fig. 5I-K). The effects followed an “inverted U” requiring optimal doses for cognitive enhancement^57^. These results underscore a fundamental, conserved role for anandamide signaling in improving working memory performance.

### AEA in thalamus increases neural activity in PFC selectively during memory periods

Working memory is known to be supported by thalamo-cortical synchrony and persistent neural representations in PFC^8–12^. How does AEA-Gpr12 mediated inhibitory signaling in thalamus facilitate excitatory responses in PFC? One possibility is that continuous cAMP depression promotes hyperpolarization mediated thalamic oscillations, that in turn support thalamocortical synchrony and persistent PFC activity. Another possibility is that inhibitory signaling in thalamus sparsifies the PFC representation thereby tuning signal to noise. To gain mechanistic insight, we delivered anandamide through intracranial perfusion locally into MD thalamus while performing longitudinal cellular resolution imaging of the PFC during working memory.

We generated a cohort of mice that were injected with CaMKIIa:GCaMP6f and implanted with a GRIN Lens targeting layers 2/3/5 in PFC, and with a pharmacology cannula above MD (Fig. S10C,D) After recovery from surgery, mice were trained on the DNMP task, and exhibited similar learning curves to earlier reported surgically naiive mice. During the testing days (30s delay period, 10 trials per day), mice were injected with either saline or AEA into MD while imaging in PFC (Methods) (Fig. 6A). We recorded a total of 739 neurons during saline sessions and 834 neurons during AEA sessions, of which we registered 332 matched neurons that were the same across both conditions (Fig. 6A, S10E,F; example matched neuron shown in 6B). Significant calcium events were extracted and aligned to behavioral events and task epochs. First, we assessed whether AEA delivery in thalamus elicited any overall changes in event rate dynamics in PFC. We found that the average event rates of all recorded neurons during the task were largely similar between both saline and AEA conditions (Fig. 6C). Next we used a session-wide permutation test to classify neurons that significantly increased activity during task-specific epochs, and found overall that the fraction of neurons tuned to each task epoch was largely similar between saline and AEA (Fig. 6D).

**Fig 6.**
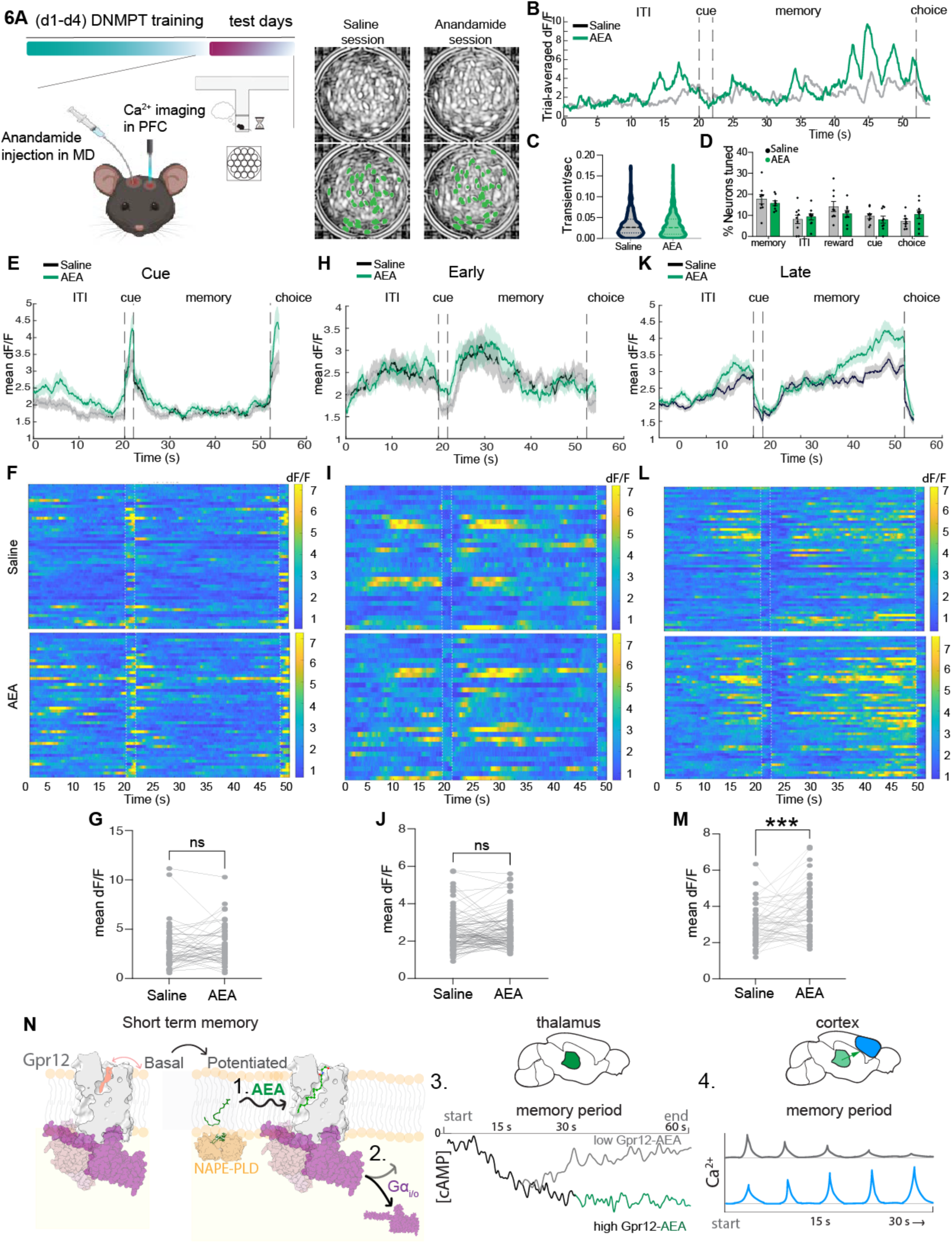
Anandamide in thalamus selectively increases neural activity in PFC during memory periods. Schematic of pharmacology in MD and imaging in PFC during DNMPT test behavior showing recording and injection sites. Mice were implanted and trained on the delay non-match-to-place task. During testing, mice received saline or anandamide (AEA) injection into MD while imaging in PFC. Right, representative field of view for PFC from one mouse, across two recording days; registered cells are labeled in green. (**B**) A representative neuron matched across AEA/Saline sessions showing the trial-averaged dF/F traces during DNMP in each phase of the task. (**C**) Distribution of average event rates during whole recording session did not differ between AEA and saline (unpaired two-tailed t-test, n=739 saline neurons and 824 AEA neurons). (**D**) Percentage of neurons tuned to task epochs under saline and AEA. Dots represent different animals. Tuning defined by exceeding 95th percentile of shuffled epoch distributions. Data are mean +/− SEM form n=9 mice. Two-way ANOVA. (**E-G**) Matched neurons tuned to the cue epoch in saline or AEA (n=49 neuron pairs) showed similar cue-period activity between conditions. **E**) population-averaged dF/F. **F**) heatmap showing individual neuron trial-averaged dF/F activity, one neuron per row, matched between saline and AEA conditions. **G**) mean dF/F did not differ between conditions; each dot is a neuron (paired two-tailed t test, n=49 neurons). (**H-J**) Matched neurons tuned to the first half of the memory epoch (Early) in saline or AEA (first 15s; n=30 neuron pairs) showed similar early memory-period activity between conditions. **H**) population-averaged dF/F. **I**) heatmap showing individual neuron trial-averaged dF/F activity, one neuron per row, matched between saline and AEA conditions. **J**) mean dF/F did not differ between neurons; each dot is a neuron (paired two-tailed t test, n=30 neurons). (**K-M**) Matched neurons tuned to the latter half of the memory epoch (Late) in saline or AEA (last 15s; n=56 neuron pairs) showed significantly higher late memory-period mean dF/F in AEA condition. **K**) population-averaged dF/F. **L**) heatmap showing individual neuron trial-averaged dF/F activity, one neuron per row, matched between saline and AEA conditions. **M**) mean dF/F was significantly higher in AEA condition; each dot is a neuron (P=0.001, paired two-tailed t test, n=56 neurons). Lines and shading indicate mean +/− SEM; heatmaps show trial-averaged dF/F for individual registered neuron pairs. Paired plots quantify epoch-mean dF/F. ns=not significant; ***=P<0.001. (**N**) Proposed model, AEA, synthesized locally by NAPE-PLD, potentiates Gpr12 in MD (1) to elicit Gi/o signaling (2), reducing cAMP during prolonged memory periods (3). This thalamic signaling cascade supports prefrontal working memory representations (4), providing a endocannabinoid-thalamocortical mechanism for working memory.

We next analyzed the dynamics of neural activity within task epochs, focusing on session-matched neurons tuned to the cue vs memory period (Fig. 6E,H,K). Plotting the mean activity of all cue-tuned neurons revealed no significant difference in activation dynamics due to AEA during the cue-period (Fig. 6E-G). However, for the memory-tuned neurons, although a similar number of neurons were recruited during the delay period for both saline and AEA conditions, we found that the mean activity levels were substantially increased by AEA (Fig. 6H-M, Fig. S10G). Interestingly, this increase was specific to neurons active in the late memory period (Fig. 6K-M; individual trials showin in Fig. 6L and quantified in Fig. 6M), corresponding to the second half of the delay period, but not to neurons active in the first-half of the memory period (Fig. 6 H-J; individual trials showin in Fig. 6I and quantified in Fig. 6J). When expanding the analysis to include all recorded cells, not just matched neurons across conditions, we again found that AEA significantly increased the mean activity of late-delay memory-tuned neurons (Fig. S10H).

Together, these results signify that anandamide does not generally increase cortical activity, but rather selectively enhnaces the persistence of neural representations associated with prolonged memory maintenance.

## DISCUSSION

Here, we identified an endogenous ligand and activation mechanism for a brain orphan receptor Gpr12. Our structural and biochemical analysis revealed that GPR12 contains two regulatory sites, a canonical orthosteric site as well as a membrane embedded side pocket, that together enable a switch from a basal to potentiated state of the receptor. *In vivo* thalamic lipidomics followed by cell-based assays identified anandamide as a potent endogenous ligand driving this potentiated state. Gpr12-containing thalamic synapses harbor the enzymatic machinery necessary for on-demand anandamide synthesis, and GPR12 activation produces sustained cAMP depressions that support the temporal window of memory maintenance. Together, these findings reveal a lipidic anandamide-Gpr12 signaling axis in thalamus that functions to extend the fidelity of memory maintenance during working memory (Fig. 6N).

While cAMP signaling has been implicated in working memory^13,14^, the real-time dynamics during behavior were unknown. Our results identify a downward ramping of cAMP levels in thalamus, that can persist up to a minute, and that matches trial-by-trial duration of memory maintenance. Despite this cAMP depression being an inhibitory signal, we find that it supports sustained neural activity in PFC, particularly at extended delays. It is interesting to consider how cAMP depressions in thalamus support neural activity persistence in PFC. Since the thalamus can intrinsically generate oscillations by converting inhibitory input into excitatory spikes, one possibility is that cAMP depressions and GIRK/HCN^58,59^ modulation work together to promote hyperpolarization mediated thalamic oscillations^60^, or shifts in tonic to bursting firing modes, that in turn support thalamocortical synchrony to prolong memory maintenance. Such a model would mechanistically link prefrontal models of working memory on short-time scales with the unique biophysical properties of the thalamus capable of supporting cortex on extended time-scales.

It is somewhat surprising that direct administration of a cannabinoid can improve working memory. It has long been appreciated that high levels of phytocannabinoids (such as THC, the primary psychoactive compound in cannabis) and endogenous cannabinoids (of which 2AG is most well studied) can diminish memory functions, with increasing doses linked to off-target inflammatory responses as well. Interestingly, both phytocannabinoids and 2AG strongly engage CB1Rs in hippocampus whereas anandamide is a comparatively weak CB1R agonist. Here, we believe the AEA-Gpr12 interaction to be potent and specific to thalamus, thus providing a targeted mechanism for working memory enhancement. Classically, endocannabinoid 2-AG functions through retrograde trans-synaptic signaling^61^. For anandamide, our data favor local synthesis and travel through the membrane to engage the side pocket of Gpr12, which has also been proposed for the AEA-TRPV1 interaction^62^. The proximal NAPE-PLD, Gpr12, FAAH1 network we observe in thalamic neurons may therefore be spatially and temporally tuned to support the rapid onset yet sustained execution of working memory during ongoing behavior. This work supports a growing view that the functions of the endocannabinoid system are shaped by the route of ligand availability and cell-type organization, enabling cannabinoid signaling to exhibit circuit specific roles across the brain^63–65^.

More broadly, with emerging studies revealing consistent lipid abnormalities across neurological diseases-including multiple sclerosis, Schizophrenia, Parkinson’s disease, and Alzheimer’s disease-each marked by memory impairments, dissection of the temporal and causal relationships between lipid dysregulation and cognition will become valuable. While classically viewed as modulators of membrane architecture^66^ and a source of energy in the nervous system^67^, the anandamide-Gpr12 interaction supports an active and dynamic signaling role for membrane lipids, suggesting broader roles for lipid signaling in higher-order cognition.

Finally, the identification of a novel membrane embedded regulatory side-pocket in Gpr12 opens new opportunities for developing compounds that modulate receptor activity with high specificity. Across class A GPCRs, drug-discovery efforts increasingly target allosteric sites, as their lower sequence conservation compared to orthosteric pockets reduces drug off-target activity^68^. Likewise, this side pocket shares low homology with family members Gpr3 and Gpr6, consistent with high regional specificity of other membrane embedded sites on GPCRs, making it particularly attractive as a druggable target. The observation that increased anandamide enhances primate working memory motivates future efforts to directly target Gpr12 as a strategy to improve cognitive function in disease and ageing.

## ACKNOWLEDGEMENTS

We thank the Rajasethupathy laboratory members for helpful discussions throughout. We thank Steve Carr, Namrata Udeshi, and Khanh Nguyen at the Broad Institute, MIT for their guidance on sample preparation and analysis of the proteomics data. We thank Corinne Lutomski and Carol Robinson for helpful discussions and analysis of some of the lipidomics experiments, as well as Agni Ghosh, Aashish Manglik, Jordan Matheisen, and Thomas Sakmar for helpful discussions related to some of the biochemical assays. We thank the core facilities at the Rockefeller University (Fisher Drug Discovery Center and particularly Chloe Larson, Cryo-Electron microscopy, Transgenic and Reproductive Technology and CRISPR and Genome Editing), at the Memorial Sloan Kettering Cancer Center (Cell Metabolism Core Facility) and at the University of Arizona (Viral Production Core). The MERFISH images in Fig 3 were adapted from the Brain Knowledge Platform, in accordance with the Allen Institute citation and rights policies, and used under their non-commercial open-access license. This work was supported by Robertson Therapeutic Development Fund and Kellen Women’s entrepreneurship Fund (A.B.O.), the Black Family Therapeutic Development Fund (A.B.O. and J.B.), the NIH Medical Scientist Training Program T32GM152349 (C.C.), the Simons Foundation (L.J.S.), HHMI (V.R.), the Halis Philanthropic Fund, the Helman Fund, Mathers and Vallee Foundations and the National Institutes of Health under award number R01MH133201 (P.R.).

## AUTHOR CONTRIBUTIONS

J.N.B. and P.R. conceived the study. J.N.B., A.B.O. and P.R. designed the experiments. N.P. performed the CryoEM experiments and related data analysis, supervised by V.R. J.N.B., A.B.O, M.K. and J.F. optimized the protocol for the cell-based assay and performed the experiments. J.N.B., M.K., J.F. and C.C. performed the surgeries. J.B and C.C. ran the proteomics experiments and C.C. developed a pipeline for the lipidomics experiments and performed sample preparation. A.B.O. ran the behavioral and photometry experiments and J.N.B. and L.J.S. analyzed the photometry data. A.L. and G.R. performed and analyzed the *in vivo* pharmacology and imaging experiments. A.F.T.A. designed and supervised the non-human primate experiments. J.N.B. and P.R. wrote the manuscript with input from all authors. P.R. supervised all aspects of the work.

## DATA AVAILABILITY

All data used in the study are available on our lab’s github page and also from the corresponding author upon request. The cryo-EM model and map will be deposited in the Protein Data Bank and Electron Microscopy Data Bank, respectively, upon paper acceptance.

## MATERIALS & METHODS

### Cryo-EM Structure & Analysis

#### Cryo-EM construct Design

The full-length human GPR12 open reading frame (ORF) was cloned under the control of a pCAG promoter. The construct contained an N-terminal HA signal peptide followed by a FLAG epitope tag and mVenus, separated from the receptor by a human rhinovirus 3C (HRV3C) protease cleavage site. The GPR12 ORF was followed by a GS12 linker (GGSS)_3_ and a C-terminal miniGs399. For Gβγ expression, human GNB1 was cloned into a single expression plasmid containing an N-terminal sfGFP tag followed by an HRV3C protease cleavage site and the full-length GNB1 ORF. An internal ribosome entry site (IRES) was included downstream of GNB1 to drive expression of tag-free human GNG2. For nanobody expression, the Nb35 coding sequence was cloned into the pET22b vector with a C-terminal glycine–serine linker followed by a FLAG epitope tag.

#### Nb35 expression and purification

Nb35 was expressed in Escherichia coli BL21 One Shot cells transformed with the Nb35 expression plasmid. Four independent colonies were picked and grown overnight at 37 °C with shaking at 200 rpm in Terrific Broth (TB). The following day, each starter culture was diluted 1:50 into 1 L TB and grown at 37 °C until reaching an OD₆₀₀ of 0.6–0.8. Protein expression was induced with 1 mM IPTG, and cultures were incubated overnight at 25 °C with shaking. Cells were harvested by centrifugation at 6,000 × g for 20 min at 4 °C, and pellets were snap-frozen in liquid nitrogen. For lysis, frozen cell pellets were thawed on ice and resuspended in SET buffer sucrose buffer supplemented with protease inhibitor (HALT) and Pierce Universal Nuclease (1:10,000 dilution). The suspension was adjusted to a total volume of 70 mL and gently stirred for 2 h at 4 °C. Osmotic shock was induced by the addition of two volumes of ice-cold Milli-Q water, followed by vigorous stirring at 4 °C. NaCl, CaCl₂, and MgCl₂ were then added to final concentrations of 150 mM, 2 mM, and 2 mM, respectively. Insoluble material was removed by ultracentrifugation at 100,000 × g for 30 min at 4 °C, and the clarified supernatant was filtered through a 5 µm PVDF filter.

The lysate was incubated overnight at 4 °C with anti-FLAG M2 agarose resin (Sigma), pre-equilibrated in extraction buffer. Resin was collected by centrifugation at 2,000 × g for 10 min at 4 °C and washed twice with 10 column volumes of wash buffer (20 mM HEPES, pH 7.5, 100 mM NaCl, 5 mM EDTA, 2 mM CaCl₂). Bound protein was eluted by incubation with 3× FLAG peptide (2 mg total) for 1 h at room temperature with end-over-end rotation. Eluted nb35 was concentrated with a 10kDA MWCO column and quantified with a 660nM pierce assay. Fractions were spiked with 10% glycerol prior to flash freezing and stored at –80c until use.

#### Expression and purification of the heterotrimeric Gpr12 complex

HEK293S GnTI⁻ cells (ATCC CRL-3022) were maintained in FreeStyle medium supplemented with 2% fetal bovine serum (FBS) and grown to a density of 1 × 10⁶ cells mL⁻¹. Cells were co-transfected with plasmids encoding Gpr12–miniGαs and Gβγ using PEI Max. Following transfection, cells were cultured for 48 h before harvesting by centrifugation at 6,000 × g for 5 min at 4 °C. Transfection efficiency was monitored by quantification of GFP-positive cells. Cell pellets were flash-frozen in liquid nitrogen and stored at −80 °C until further use.

Frozen pellets were thawed on ice and resuspended in three volumes of extraction buffer at 4 °C with end-over-end rotation until fully dispersed, followed by stirring for 1 h at 4 °C. The lysate was clarified by ultracentrifugation at 100,000 × g for 30 min at 4 °C, and the supernatant was filtered through a 5 µm PVDF membrane. The clarified lysate was incubated for 2 h at 4 °C with custom-generated GFP nanobody agarose resin (5 mL of a 50% slurry), pre-equilibrated in extraction buffer, with gentle stirring. Resin was collected using a Bio-Rad 50 mL gravity-flow column, and bound complexes were released by incubation with HRV3C protease (1 mg) and Nb35 (200 µg) for 1 h at 4 °C. Eluted protein was concentrated using a 100 kDa molecular weight cutoff concentrator, filtered through a 2 µm membrane, and subjected to size-exclusion chromatography by HPLC. Peak fractions were analyzed by rapid Coomassie staining, pooled, and concentrated to ∼5 mg mL⁻¹ prior to grid preparation. Cryo-EM grids were prepared using Quantifoil Au R1.2/1.3 400 mesh and UltraFoil R1.2/1.3 300 mesh grids. Grids were plasma cleaned prior to sample application. Using a Vitrobot, 3 µL of protein sample was applied to each grid with a 30 second wait time followed by blotting for 1-3 seconds before plunge-freezing in liquid ethane.

#### Cryo-EM data processing and model refinement

A total of 35,923 micrographs were collected over three days from two independent grids on a FEI Titan Krios transmission electron microscope equipped with a Gatan K3 direct electron detector and a Cs corrector, operated at a calibrated physical pixel size of 0.86 Å. Movie frames were motion-corrected using MotionCor2, and contrast transfer function (CTF) parameters were estimated using CTFFIND4, as implemented in RELION v5.0b3. Particle picking was performed using Topaz, followed by particle extraction in RELION. Subsequent rounds of two-dimensional classification, three-dimensional refinement, and Bayesian particle polishing were carried out in RELION v5.0b3. Selected particle stacks were further refined using non-uniform refinement in cryoSPARC v4.7.1 to improve map quality and angular sampling. Focused refinements were performed in cryoSPARC v4.7.1 using local refinement with soft masks generated from atomic model chains to improve density quality in specific regions of interest. Density modification and composite map generation were carried out using Resolve Cryo-EM in PHENIX v1.21.2-5419, and focused maps were combined to generate a final composite reconstruction. Atomic model building and refinement were performed iteratively using Coot v0.9.8.92, PHENIX real-space refinement (v1.21.2-5419), and ISOLDE v1.4, with visualization and validation conducted in ChimeraX v1.4.

### Cell-based cAMP accumulation assays

To assess the effect of point mutations on Gpr12-mediated cAMP accumulation, we performed transient transfections of the WT or Mutant Gpr12 in HEK cells and measured cAMP accumulation using the FRET based Gs HiRange assay as described below. In brief, we seeded HEK293T cells into 384-well white plates with transparent bottoms (Greiner Bio-One, cat. no. 781098) at 5,000 cells per well. At approximately 50% confluency, cells were transfected using Lipofectamine 3000 (Thermo Fisher Scientific, cat. no. L3000008) according to the manufacturer’s instructions. Briefly, for each well, 0.075 μL of Lipofectamine 3000 was diluted in 2.5 μL of Opti-MEM, and 50 ng of plasmid DNA was diluted in 2.5 μL of Opti-MEM with 0.1 μL of P3000 reagent. The diluted DNA and Lipofectamine solutions were combined and incubated for 15 min at room temperature before addition to wells. Each well received a total of 5 μL Opti-MEM, 50 ng plasmid DNA, 0.1 μL P3000, and 0.075 μL Lipofectamine 3000 reagent. Twenty-four hours post-transfection, transfection efficiency was verified by GFP fluorescence from the pINDUCER21 backbone. Gpr12 expression was induced with 0.1 μg/mL doxycycline (MedChemExpress, cat. no. HY-N0565B) for 3 h. Media was then replaced with 10 μL of either media containing DMSO and 200 μM IBMX (MedChemExpress, cat. no. HY-12318) or media containing 100 μM AEA (Enzo Life Sciences, cat. no. BML-FA017) and 200 μM IBMX. Plates were sealed and incubated at 37 °C for 15 min. cAMP levels were measured using the cAMP Gs HiRange assay (Revvity, cat. no. 62AM6PEB) according to the manufacturer’s protocol. Briefly, 5 μL of cAMP-d2 reagent followed by 5 μL of Eu-cryptate antibody (both diluted in 1× lysis/detection buffer) were added to each well. Plates were sealed and incubated for 1 h at room temperature in the dark. Opaque white seals (Revvity, cat. no. 6005199) were applied to plate bottoms, and time-resolved fluorescence was measured using a BioTek Synergy Neo2 plate reader following Revvity’s imaging instructions.

To quantify Gpr12-dependent cAMP accumulation in response to various small molecules, we required a more controlled Gpr12 expression system. For these experiments we generated a doxycycline inducible HEK293 cell line expressing human GPR12. Wild-type human GPR12 was C-terminally tagged with a 2×FLAG epitope and cloned into the pcDNA™5/FRT/TO inducible expression vector (Invitrogen, V6520-20). Expression cassettes were site-specifically integrated into the genome of Flp-In™ T-REx™ 293 cells (Invitrogen, R78007) using Flp recombinase–mediated integration, yielding isogenic, single-copy inducible cell lines. Cells were maintained under standard culture conditions and split into into clear-bottom T25 flasks at a density of 0.7 x 10^6^ cells. After 48 h of growth at 37 °C with 5% CO₂, receptor expression was induced by addition of doxycycline (1 µg mL⁻¹) for 3 hours. Cells were then gently trypsinized and while in suspension diluted to a concentration of 2,000 cells/ 5uL. 5uL of cells were transferred to pre-warmed opaque white 384 well plates containing either 5uL IBMX (0.1mM), 5uL forskolin (10uM), 5uL of drug concentration, or 5uL of matched DMSO concentration. Wells were gently pipetted up and down (10uL final volume) prior to sealing and incubation for the 15 minutes at 37c. Finally, intracellular cAMP concentrations were quantified using a FRET-based HTRF cAMP Gs HiRange assay (Cisbio, 62AM6PEB) according to the manufacturer’s instructions^28^. Cell dilution, doxycycline concentration, and expression timepoints were empirically calibrated to be within the linear dynamic range of the cAMP kit to allow for absolute quantification of cAMP concentration. Fluorescence measurements were acquired on a Synergy™ Neo2 multimode plate reader (BioTek) using excitation, emission, and signal-attenuation settings recommended by the manufacturer for HTRF detection. Absolute cAMP concentrations were calculated from standard curves generated in parallel.

#### Flow cytometry for cell surface expression of Gpr12 mutants

HEK293T cells were seeded into 6 well plates at a seeding density of 0.3*10^6^ cells per well. 24 h later, cells were transfected with the hGpr12 mutant plasmid constructs with Lipofectamine 3000 according to the manufacturer protocol, with three replicate wells included per mutant, including a non-transfected control (GFP-). After 24h, successful transfection was assessed by observing the GFP+ cells under a fluorescent microscope. At this point, doxycycline (1µg/µl) was added to the media to induce expression of the hGp12-FLAG-tag protein over the course of 24h. The following day, cells were harvested with trypsin for flow cytometry, spun down 300g for 5 min, and resuspended in 700µl of FACS buffer (PBS, 0.5% w/v BSA, 0.1% w/v sodium azide). 100µl of a GFP+ sample was removed for the non-stained control (GFP-, PE-). The cells were blocked in 5% BSA for 15 min, followed by primary antibody staining (anti-Flag-PE, Biolegend cat no. 637309, 1:200) for 30 min on ice. Cells were washed 2x FACS buffer with 350g, 5 min centrifuge spins in between. Final samples were resuspended to a final volume of 400µl FACS buffer with 1µg/ml DAPI. For gating controls, GFP-PE-(no transfection) and GFP+PE-(transfection, no staining) samples were used. Flow cytometry was performed using a BD LSR-Fortessa (BD FACSDiva Software, v8.0.1) with a 100-µm nozzle. The cell suspensions were first gated on forward scatter, then within this population based on DAPI-positive cells. The cell surface expression was estimated based on the number of GFP+PE+ out of all GFP+ cells (of all transfected cells, the percentage that had cell surface FLAG-tag expression).

### Microscale thermophoresis (MST) ligand binding assay

Purified Gpr12 protein was prepared as described for cryo-EM sample preparation, except without a C-terminal miniGs fusion and without co-expression of Gβ1/Gγ2 components in HEK293S GnTI⁻ cells (ATCC CRL-3022). Immunopurified Gpr12 was concentrated, injected onto an ÄKTA micro FSEC system, and peak fractions were pooled, concentrated, and flash-frozen at 5 mg/mL. Gpr12 was fluorescently labeled using the NanoTemper RED-NHS 2nd Generation Protein Labeling Kit (Cat# MO-L011) according to the manufacturer’s instructions. Briefly, Gpr12 was buffer-exchanged from Tris-containing buffer into HEPES buffer prior to labeling, yielding a calculated degree of labeling (DOL) of 0.9.

For MST binding assays, NanoTemper Monolith Premium Capillaries (Cat# MO-K025) were loaded with 12 µL samples containing 20 nM labeled Gpr12 protein and serial dilutions of ligand. DMSO was maintained at 1% across all experiments. Anandamide and anandamide phosphate were dissolved in argon-purged DMSO prior to dilution into final assay buffer containing 50 mM HEPES pH 8.0, 100 mM NaCl, 100 mM KCl, 0.02% LMNG, 0.004% CHS, 1 mM CaCl₂, and 1 mM MgCl₂. Anandamide phosphate was included as a structurally related non-binding control. Data were collected on a NanoTemper Monolith instrument and analyzed using MO.Affinity Analysis 3 software. Experimentally derived MST ratios were plotted against ligand concentration, and Kd values were calculated using a quadratic binding function within the MO.Affinity Analysis 3 software.

### Lipidomics

#### Tissue extraction

For targeted lipidomics experiments, two cohorts of mice were injected in the mediodorsal thalamus (see Surgeries) with AAV-Gpr12-V5-mCherry or AAV-V5-mCherry (controls). The viruses were expressed for three weeks before animals were euthanized. Animals were perfused with 1x PBS, and the mediodorsal thalamus was micro-dissected under fluorescence guidance using a handheld fluorescent flashlight to visualize the targeted area. Tissue was snap-frozen on liquid N_2_ until tissue processing. For processing, 500 µl of lysis buffer (1% LMNG, 1% DDM, 1% GDN, 100 mM NaCl, 100 mM KCl, 50 mM Tris pH 8.0, 1 mM CaCl₂, 1 mM MgCl₂, Halt protease inhibitor [1x], and 20 µg ml⁻¹ DNase) was added to each tissue sample (one mouse thalamus per sample). The tissue was broken up by pipetting up and down through a P1000 tip, followed by further dissociation by running the lysate through a 23G, then 27G syringe. The samples were kept on ice for 30 minutes with light resuspension every ten minutes through the 27G syringe. Samples were spun down 15,000 x g for 20 min at 4°C in a swing bucket rotor tabletop centrifuge. The supernatant was removed and transferred to a new tube. 1.4 mL 0.1% detergent lysis buffer (0.1% LMNG, 0.1% DDM, 0.1% GDN, 100 mM NaCl, 100 mM KCl, 50 mM Tris pH 8.0, 1 mM CaCl₂, 1 mM MgCl₂, Halt protease inhibitor [1x], and 20 µg ml⁻¹ DNase) was added to the 0.5 mL lysate. After washing magnetic V5 agarose beads (ChromoTek) 3x with 1 mL 0.1% detergent lysis buffer, 25µl of the beads was transferred into each lysate sample for bead incubation. 50µl of the input lysate (input fraction) was saved. Beads were incubated in the lysate with end-over-end rotation at RT for 1h. The flowthrough (flowthrough fraction) was collected and saved for downstream analysis. The beads were washed 3x by resuspending in 0.1% detergent lysis buffer. On the last wash, beads were transferred to a new tube, and the remaining buffer was removed. To elute the V5-tagged protein on the beads, 100µl of 1 mg/mL V5 peptide (ChromoTek) was added to the beads and resuspended with a pipette. Beads were incubated in the V5 peptide with end-over-end rotation at RT for 10 min. The supernatant collected, and the elution was repeated for 2-3 more times. The pooled supernatants were spun down in a Pierce Concentrator, PES, 10K MWCO 0.5 mL column (ThermoFisher) to concentrate the sample, spinning down in a tabletop centrifuge at 15,000 x g for 15 min at 4°C. The concentrated sample (V5 elution fraction) was transferred into a new tube and snap frozen on liquid N_2_ for downstream lipidomics processing. To assess the leftover protein bound to the beads, a harsh elution as performed by boiling the beads in 2X SDS Laemmli buffer at 95°C, collecting the supernatant (SDS fraction). For western blot analysis of the immunoprecipitation (IP) preparation, the following fractions were used: input, flowthrough, concentrated V5 elution, SDS fractions. For V5 western blot analysis, a western blot was performed as previously described (Methods), with V5 (Invitrogen, cat no. R960-25, 1:1,000 dilution) and IRDye 800CW Goat anti-Mouse IgG Secondary Antibody (Li Cor Biotech, cat no. NC99401841). Full, uncropped western blot images are provided in Supplementary figures.

#### Lipid extraction

Approximately 30 mg of tissue was placed into a bead-beating tube and homogenized in 60:40 (v/v) methanol:water at a ratio of 1:15 (w/v), supplemented with internal standards (d31-palmitate and SPLASH Lipidomix; Avanti Polar Lipids, cat. no. 330707), using a BeadRuptor homogenizer. Following homogenization, 400 µL of lysate was transferred to a new microcentrifuge tube, mixed with 480 µL chloroform, vortexed, and incubated on ice for 10 min. Samples were then centrifuged at 20,000 × g for 5 min to achieve phase separation. The lower organic phase (450 µL) was collected using a Hamilton syringe and transferred to a glass vial. A second extraction was performed by adding 450 µL chloroform to the remaining aqueous phase, followed by vortexing, incubation on ice for 10 min, and centrifugation at 20,000 × g for 5 min. The organic phases were combined, split into two aliquots, and dried under nitrogen. Derivatization of anandamide and arachidonic acid was performed using a optimized protocol^69^. Dried extracts were resuspended in 60 µL acetonitrile, followed by sequential addition of 20 µL HATU (50 mM) and 20 µL diethylamine (DEED; 100 mM), with vortexing between additions. Samples were incubated at room temperature for 1 min, centrifuged at maximum speed for 20 min, and 90 µL of the supernatant was transferred to glass vials with conical inserts for LC-MS analysis. For untargeted lipidomic analysis, dried extracts were resuspended in 50 µL of acetonitrile:isopropanol (1:1, v/v), vortexed, and incubated at room temperature for 20 min. Samples were centrifuged at 20,000 × g for 20 min, and the supernatant was transferred to 40 µL glass inserts in glass vials for LC-MS analysis.

#### Lipid analysis

Prior to experimental runs, a diagnostic system suitability test (SST) consisting in a SPLASH lipid mixture was run on the Agilent 6546 LC/Q-TOF LC-MS system to verify instrument performance. Individual species were reviewed for peak shape, chromatographic resolution, and stability/mass accuracy over time. Brain tissue samples were normalized by mass and subjected to a two-phase organic extraction prior to analysis, with deuterated internal standards added before extraction to control for recovery and technical variability. Anandamide and arachidonic acid peak areas were empirically quantified using calibration runs with known concentrations. Consistent extraction efficiency was confirmed by low variability of internal standard signals across samples (CV < 10%). Retention time reproducibility was monitored across pooled QC injections. Anandamide and arachidonic acid peak areas were empirically quantified using calibration runs of known concentrations and normalized to the spiked in d31 palmitate internal standard during targeted analysis.

Lipid feature detection and annotation were established through iterative injections of pooled quality control samples interspersed through the acquisition sequence. Tandem mass spectrometry (MS/MS) spectra were acquired on the Aglient 6546 LC/Q-TOF using quadrupole precursor isolation followed by collision-induced dissociation, enabling high resolution fragmentation analysis for structural validation of lipid species. MS/MS spectra were assessed against reference fragmentation pattern. Lipidmaps, LipidCruncher, LipidEx, and SIRIUS were used for lipid species analysis. Peak picking and integration were performed using Skyline-Daily v.25. After initial peak picking and integration, additional stringent data filtration steps were applied such as removal of compounds with coefficient of variation (CV) greater than 30% in pooled quality control runs in addition to removal of species with high signal during blank runs. Individual lipid species were first normalized to the peak area of their corresponding deuterated internal standard prior to statistical analysis. Filtered abundances were subsequently used for species-level comparison of log_2_ abundance using to-tailed student’s t-tests.

### Proteomics

#### TurboID Construct Design & Surgeries

For TurboID experiments, six-week old wildtype C57BL/6 mice were injected with 1µl AAV-Gpr12:TurboID, AAV-nuclear export signal (NES):TurboID, AAV-β2ar:TurboID at [1 x 10^13^ GC/mL], bilaterally in the mediodorsal thalamus (A/P –1.35, M/L +/− 0.4, D/V –3.4). For steady state experiments, the incisions were closed with 4-0 vicryl sutures post-surgery until re-opening for surgery at the time of biotin injection. For behavioral experiments mice were implanted with a unilateral custom made optofluid cannula (Doric; M3, 250µm guiding tube width, 2.3mm depth) at the coordinate A/P –0.5, M/L +1.0, D/V –2.3 to provide chronic access to the lateral ventricle. mice used for head-fixed behavior, a custom titanium headplate was adhered to the skull with Metabond. For mice that required cannulas, the cannulas were implanted immediately following viral injection. The TurboID viruses were allowed to express for 5 weeks for optimal viral expression. Steady state animals underwent surgery and direct injection of 10mM biotin (Sigma Aldrich #B4640); PBS) into the lateral ventricle at the time of biotin labeling. For awake animals with implanted cannulas, they were head-fixed for the duration of the biotin injection. The optofluid cannulas were fit to a fluid injector (Doric, M3 barrel) and 25G tubing (Doric) and connected to a [pump system] for slow, controlled injections at a rate of 100nL/min and a volume of 400nL. The biotin labeling window for all experiments was 2 h. For behavioral experiments, the trials were run in the last 30 minutes of the 2 h window. At the end of the biotin labeling window, animals were deeply anesthetized before perfusion with 4°C PBS to slow the enzymatic activity of the biotin ligase. The brain was rapidly dissected and specific region of interest micro-dissected. Samples were snap frozen with liquid N_2_ and stored at –80°C until the time of tissue processing.

#### Whole Neuron Purification

Lysis buffers were prepared as follows. A stock lysis buffer containing 50 mM Tris-HCl (pH 7.5), 150 mM NaCl, 1% (w/v) SDS, 0.25% (w/v) sodium deoxycholate and 1% (v/v) Triton X-100 in deionized water was first prepared. To 30 ml of this buffer, 59 mg sodium L-ascorbate, 38 mg Trolox and 20 mg sodium azide were added. From this modified buffer, 20 ml was supplemented with Halt protease inhibitor cocktail (1x) and 200 µl 1 mM PMSF to generate the Active Lysis Buffer (ALB). Tissue samples were thawed on ice and homogenized sequentially in ALB: initially, 200 µL was added and tissues were minced using spring scissors. An additional 300 µL was added, and the homogenate was passed through a 28G insulin syringe five times. This was followed by the addition of 500 µL ALB and three additional passes through the syringe. A final 400 µL was added to reach a total volume of 1.4 mL per sample. Samples were kept on ice until all were processed. Lysates were sonicated (6 cycles of 5 seconds on, 10 seconds off) and cleared by centrifugation at 15,000 × g for 5 minutes at 4 °C. The supernatant was collected and denatured at 95 °C for 5 minutes, followed by incubation on ice for 10 minutes. A second centrifugation was performed at 20,000 × g for 15 minutes at 4 °C, and the clarified supernatant was collected for downstream applications.

#### Enrichment of biotinylated proteins

Biotinylated proteins were purified using streptavidin-conjugated magnetic beads. Briefly, 50 µL of beads per sample were washed twice with 1 mL ALB. Clarified lysates were incubated with the beads overnight at 4 °C on a rotator. On the following day, buffers for washing were freshly prepared. Sodium carbonate solution (0.1 M) was prepared by dissolving 529 mg Na₂CO₃ in 50 mL sterile H₂O. Urea buffer (2 M urea in 10 mM Tris-HCl) was made by mixing 2.5 mL of 8 M urea and 100 µL of 1 M Tris-HCl, and bringing the volume to 10 mL with sterile H₂O. KCl wash buffer (1 M) was prepared by dissolving 746 mg KCl in 10 mL sterile H₂O. After overnight incubation, the flow-through was collected, and the beads were sequentially washed: RIPA 2 min (2X), 1M KCl 2 min, 0.1M Na_2_Co_3_ 30s, 2M Urea in 10mM Tris-Cl 30s, RIPA 2min (3X). On the third RIPA wash, beads were transferred to a new tube. Samples were kept on ice until further processing.

#### Elution of biotinylated proteins

Elution buffer was prepared by first generating a 6× sample buffer (6× SB) consisting of 16.5 mL 1 M Tris-HCl (pH 6.81), 17 mL glycerol, 4.5 g SDS, and 45 mg DTT, brought to a final volume of 50 mL with molecular-grade H₂O. For each elution, 4 mL of elution buffer was made fresh by mixing 2 mL 6× SB, 1,840 µL H₂O, 80 µL 100 mM biotin, 80 µL 1 M DTT, and 20.86 mg TUDCA. Beads were pelleted using a magnetic rack and resuspended in 50 µL of elution buffer. Samples were incubated at 95 °C for 10 minutes and allowed to cool at room temperature for 5 minutes before being placed on a magnetic rack. The supernatant containing the eluted proteins was transferred to fresh tubes and stored at – 20 °C until further analysis.

#### On-bead trypsin digestion of biotinylated proteins

For each sample, biotinylated proteins bound to streptavidin magnetic beads were washed four times with 200 uL of 50 mM Tris-HCl (pH=7.5) buffer. The final wash was then removed, and 80uL of the digestion buffer (2M Urea, 50nM Tris-HCl, 1mM DTT, and 0.4ug trypsin) was added for incubation at room temperature (RT) while shaking at 1000 rpm. Incubation was conducted for 1 hour, and the supernatant was collected and transferred into a separate tube. This process was then repeated for another round with 30-minute incubation. After the second supernatant was collected, 60 uL of 2M Urea/50mM Tris-HCl buffer was added to the tube for 2x washes, which were collected and pooled with the digestion supernatant. The pooled eluate was spun down at 5000 x g for 30 sec. Next, the supernatant was collected and reduced with 4 mM DTT for 30 minutes, followed by alkylation with 10 mM Iodoacetamide for 45 min in the dark at RT while shaking at 1000 rpm. Finally, each sample was further digested overnight with 0.5ug of trypsin on a Thermo shaker at RT and 1000rpm shaking.

The following morning, neat formic acid (FA) was used to acidify digested peptide samples to the final concentration of 1% FA (pH<3). Samples were then desalted using in-house packed C18 (3M) StageTips. Briefly, C18 StageTips were conditioned sequentially with 100 uL of 100% methanol (MeOH), 100 uL of 50% (vol/vol) acetonitrile (MeCN) with 0.1% (vol/vol) FA, and 2x 100uL of 0.1% (vol/vol) FA. Next, acidified peptides were loaded onto the C18 StageTips and washed twice with 100 uL of 0.1% FA. Desalted peptides were then eluted from the C18 resin using 50uL of 50% MeCN/0.1%FA, snap-frozen, vacuum-centrifuged until dry.

#### TMT labeling and StageTip peptide fractionation

Each peptide sample was resuspended in 80 μL of 50 mM HEPES and labeled with 20 uL of the 25 ug/uL TMTpro reagents in MeCN (Thermo Fisher Scientific). Samples were then incubated 1 hour at RT while shaking at 1000 rpm. TMT-labeling reaction was quenched by incubating with 4 μL of 5% hydroxylamine for 15 min at RT with 1000rpm shaking. Next, TMT-labeled peptides from all samples were pooled into one tube and vacuum-centrifuged to dry. The samples were then reconstituted in 200 μL of 0.1% FA, desalted on C18 StageTips, and dried to completion.

The labeled peptides underwent basic reverse phase (bRP) fractionation on an in-house packed SDB-RPS (3M) StageTip. Specifically, the SDB-RPS StageTip was conditioned sequentially with 100 μL of 100% MeOH, 100 μL 50% MeCN/0.1% FA, and 2x 100 μL 0.1% FA. Peptides were resuspended in 200 uL of 0.1% FA, loaded on the conditioned StageTip, and then eluted from the StageTip in eight fractions using 20 mM ammonium formate buffers with increasing (vol/vol) concentrations of MeCN (5%, 7.5%, 10%, 12.5%, 15%, 20%, 25%, and 45%). The eight fractions were then vacuum-centrifuged until completely dry.

#### Liquid chromatography and mass spectrometry

Peptide samples were analyzed on an online liquid chromatography tandem mass spectrometry (LC-MS/MS) system, including a Vanquish Neo UPHLC (Thermo Fisher Scientific) coupled to an Orbitrap Exploris 480 (Thermo Fisher Scientific). Previously collected eight fractions were reconstituted in 9 uL of 3% MeCN/ 0.1% FA, and 4 uL of each fraction was injected onto an in-house packed microcapillary column (Picofrit with 10 µm tip opening/75 µm diameter, New Objective, PF360-75-10-N-5) with 30 cm of C18 silica material (1.5 µm ReproSil-Pur C18-AQ medium, Dr. Maisch GmbH, r119.aq), heated to 50 °C using column heater sleeves (PhoenixST). Peptides were eluted into the Orbitrap Exploris 480 at a flow rate of 200 nL/min. Each fraction was run on a 110min-method, including a linear 84 min gradient from 94.6% solvent A (0.1% formic acid) to 27% solvent B (99.9% acetonitrile, 0.1% formic acid), followed by a linear 9 min gradient from 27% solvent B to 54% solvent B.

Mass spectrometry was conducted in a data-dependent acquisition mode. MS1 spectra were measured with 60,000 resolution, 300% normalized AGC target, and m/z range from 350 to 1800. MS2 spectra were acquired for the top 20 most abundant ions per cycle at 45,000 resolution, 30% AGC target, 0.7 m/z isolation window, and 34 normalized collision energy. The dynamic exclusion time was set to 20 s, and the peptide match and isotope exclusion functions were enabled.

#### Mass spectrometry data processing

Mass spectrometry data were processed using Spectrum Mill Rev BI.07.11.216 (proteomics.broadinstitute.org). Raw file extraction retained spectra within a precursor mass range of 600-6000 Da with a minimum MS1 signal-to-noise ratio of 25. Additionally, MS1 spectra within a retention time range of +/− 45 s, or within a precursor m/z tolerance of +/− 1.4 m/z were merged. MS/MS searching was performed against a human Uniprot mouse database, released on April 07, 2021. Digestion parameters were set to “trypsin allow P” with an allowance of 4 missed cleavages. The MS/MS search included fixed modifications, carbamidomethylation on cysteine and TMTPro on the N-terminus and internal lysine, and variable modifications, acetylation of the protein N-terminus, oxidation of methionine, N-term Q-pyroglutamate formation, and N-term deamidation. The matching tolerances were set with a minimum matched peak intensity of 30%, precursor and product mass tolerance of +/− 20 ppm, and a maximum ambiguous precursor charge of 3. Peptide spectrum matches (PSMs) were validated with a maximum false discovery rate (FDR) threshold of 1.2% for precursor charges ranging from +2 to +6. A target protein score of 0 was applied during protein polishing autovalidation to further filter PSMs. TMTpro reporter ion intensities were corrected for isotopic impurities using the afRICA correction method in the Spectrum Mill protein/peptide summary module, which utilizes determinant calculations according to Cramer’s Rule. Protein quantification and statistical analysis were performed using the Proteomics Toolset for Integrative Data Analysis (Protigy, v1.0.7, Broad Institute, https://github.com/broadinstitute/protigy). Differential protein expression was evaluated using moderated t-tests, with P-values calculated to assess significance.

Each protein was associated with a log2-transformed ratio of every TMT condition to the median intensity of all channels. Protein data were then normalized to the median within each condition group. After normalization, an empirical Bayes-moderated t-test was used to compare treatment groups using the limma R package. P-values associated with every modified peptide were adjusted using Benjamini–Hochberg FDR approach.

#### Western blotting

Magnetic beads were pelleted and 5 µL of remaining condensation was added back to each sample. For gel loading, samples were prepared with 2.5 µL 6× SB, 2.5 µL H₂O, and 10 µL of sample. Gels were run at 150 V under standard conditions. Following electrophoresis, PVDF membranes were soaked in 100% methanol, equilibrated in H₂O for 1– 2 minutes, and transferred to Towbin transfer buffer (without SDS). Gels were soaked in transfer buffer for 10 minutes before transfer. Proteins were transferred at 70 V for 1 hour at 4 °C. After transfer, membranes were incubated in 100% methanol for 1 minute, rinsed with Milli-Q H₂O, and equilibrated in PBS for 2 minutes. Membranes were blocked for 1 hour at room temperature in 5% BSA prepared in PBS-T (PBS with 0.1% Tween-20). Primary antibody incubation was performed overnight at 4 °C in 20 mL blocking buffer supplemented with 20 µL Tween-20 and 10 µL rabbit anti-HA antibody (Cell Signaling Technology, cat no. 3724, 1:2,000 dilution). After four 5-minute washes with PBS-T, membranes were incubated with secondary antibodies for 1 hour at room temperature in 20 mL blocking buffer containing 20 µL Tween-20, 20 µL 10% SDS, 1 µL goat anti-rabbit IRDye 800CW (Licor, cat no. 926-32211, 1:20,000 dilution), and 4 µL streptavidin IRDye 680RD (Licor, cat no. 926-68079, 1:5,000 dilution). Membranes were then washed three times for 5 minutes in PBS-T and once in PBS for 5 minutes prior to imaging.

### Mice

All mice used in this study were C57Bl6/J purchased from the Jackson Laboratory. Male and Female mice were separately group housed three to five per cage and kept under a 12 h light– dark cycle in a temperature-controlled environment with ad libitum food and water unless animals were food restricted for behavioral testing. Animal ages Six to Eight weeks were used for Stereotaxic Viral Injections and Cannula implantations with behavioral testing performed from 10 to 12 weeks aged. All experiments involved mixed sex cohorts with number of mice used for each experiment determined from expected variance and effect size from previous studies. No statistical methodology was used to predetermine sample size. All procedures were done in accordance with pre-approved guidelines set by the Institutional Animal Care and Use Committee (IACUC) (protocol no. #22087-H) at the Rockefeller University.

### Generation of Gpr12-3xHA knock-in mice

A c-terminal 3xHA epitope tax was introduced in frame to the endogenous mouse *Gpr12* locus using standard CRISPR-cas9 mediated genome editing in a C57BL/6J background. Briefly, two guide RNAs targeting the c-terminus of *Gpr12* at the end of exon 2 (Gpr12cKO-gRNA-G: 5′-GGTAGAGGCAGCGTCCTATG-3′; Gpr12cKO-gRNA-H: 5′-GGCAGCGTCCTATGAGGAGA-3′) and a single stranded DNA donor encoding the 3xHA sequence with flanking homology arms to *Gpr12* were microinjected with Cas9 mRNA into one cell embryos. Injected embryos were then transferred into pseudopregnant mice. Founder mice were genotyped using PCR primers flanking the edited locus Gpr12-SA-F5: 5′-CCTTGATCGCCGATTACACC-3′; Gpr12-SA-R6: 5′-TGGCAAAGCTCCTATCCTGT-3′), yielding a 650-bp wild-type product and a 739-bp 3xHA knock-in product, followed by ZraI restriction digest, which selectively cleaves the edited allele (476 bp and 263 bp). Allelic composition and mosaicism were further assessed by Sanger sequencing and DECODR analysis. Seven founders carrying the 3xHA insertion were identified, exhibiting variable degrees of mosaicism; founders with higher proportions of intact 3xHA alleles were prioritized for germline transmission. Prior to experimental use, Gpr12-3xHA mice were compared with wild-type littermates and exhibited no detectable differences in body weight, baseline learning performance, or gross cognitive behavior.

### Generation of Gpr12(−/−) knockout mice

A Gpr12 knockout mouse line (m7-c98) was generated through removing a 98bp segment at the 5’ end of the *Gpr12* locus using standard CRISPR-cas9 mediated genome editing in a C57BL/6J background. Two guide RNAs spanning a 105bp region at the beginning of Gpr12 coding sequence (gRNA-A: 5′-CCTCGGGACTGTATAGATGCCGG-3′; gRNA-B: 5′-CCGAGCTCGTTGTCAACCCCTGG-3′) were used to induce a targeted deletion. Cas9 mRNA and both gide RNAs were co microinjected into one cell embryos which were subsequently transferred into pseudopregnant mice. Founder animals were screened by PCR using primers flanking the gRNA annealing sites Gpr12-SA-F2: 5′-CAGTTGTTGGTTTTCGGCATG-3′; Gpr12-SA-R1: 5′-GATGGCAGCGTTGTTCTTAGT-3′) and analyzed by sanger sequencing. A founder line carrying a 98-bp deletion was identified, resulting in a frameshift and introduction of a premature stop codon after 27 amino acids. Founder with confirmed frameshift allele as bred to establish germline transmission of the Gpr12 knockout allele and deletion was verified in subsequent generations.

### Surgeries

#### Viral injections

All surgical procedures were performed under protocols approved by the Rockefeller University IACUC. One day prior to surgery mice were provided with Meloxicam tablets. Mice were anesthetized with 2% Isoflurane and placed on a heating pad under a Kopf stereotactic apparatus with paralube vet ointment applied to eyes. Meloxicam was administered 0.2mg/kg via intraperitoneal injection in addition to 0.5mL bacteriostatic saline. Viruses were injected using a 35G beveled needle in a 10uL NanoFil injection syringe (World Precision Instruments) using a nanoliter injection pump (Harvard Apparatus). Prior to injection, needles were placed at injection site for 2 minutes. All viruses were injected at a rate of 100nL min^-1^. After injection, the needle was raised 0.1mm and allowed to sit for 5 minutes to prevent backflow of the virus. For Gpr12 Overexpression experiments, 1000nL (unless otherwise mentioned) of AAV9 packaged pHsyn-Gpr12-V5-IRES-TdTomato (custom plasmid packaged by University of Arizona Viral Packaging Core) at a titer of 1 x 10^13^ GC/mL was delivered bilaterally to the mediodorsal thalamus (A/P –1.25mm, D/V –3.4mm, M/L +-0.4mm). For GFLAMP2 recordings, 1000nL of 5 x 10^12^ GC/mL of AAV9 packaged camKIIa-GFLAMP2 (custom plasmid packaged by University of Arizona Viral Packaging Core) was delivered unilaterally in mdTH (A/P –1.25mm, D/V –3.4mm, M/L –0.4mm).In the case of multiple viruses injected in the same location (i.e., Gpr12 WT/Mutant/mCherry + GFlamp2), viral stocks were mixed at the given final titer and delivered in a single injection. pHsyn-mcherry packaged in AAV9 (Addgene cat#114472-AAV9) was injected at the same titer (1 x 10^13^ GC/mL) as the Gpr12 Overexpression plasmid.

#### Implanting fiberoptic cannulas

After standard anesthesia as above, animals were additionally administered intramuscular dexamethasone (0.2 mg kg^-1^) and maintained under anesthesia at 1.5-2% for duration of procedure. An anterior-posterior incision was made and the skin directly overlaying implantation sites is removed to allow for custom 10mm titanium headplate to affix to the skull at the end of the procedure. Fiber optic implantation occurred directly after all viral injections on a given animal were completed. Mice were implanted with 1.25mm ferrule-coupled optical fibers (0.48 NA, 400µm diameter, Doric Lenses) cut to a custom length so that the implantation site was 0.2mm dorsal to the viral injection site. Cannula implants were slowly lowered using a stereotaxic cannula holder (Doric) at a rate of 1mm/min. Optical Glue (Edmund Optics) was used to stable affix cannulas in position until all were completed. Finally, a custom 10mm diameter titanium headplate was placed on the skull, flush with the surrounding skull skin, and the headplates and optical cannulas were securely fixed using Metabond adhesive cement.

#### Preparation of Anandamide and solutions

Anandamide was purchased as a neat oil (Enzo Life Sciences, BML-FA017) and stored at – 80°C under inert conditions until use. Polyunsaturated lipids, including anandamide and arachidonic acid, are susceptible to spontaneous oxidative degradation driven by light, oxygen exposure, and heat. Although methanol or chloroform-based solvents are sometimes used for lipid storage, DMSO was selected here due to its high solubility and its compatibility with *in vivo* and cell-based applications. To minimize peroxidation, anandamide stocks were prepared immediately prior to experiments under an argon stream using freshly argon purged anhydrous DMSO. Anandamide was dissolved to 50mg/mL, and complete solubilization was confirmed by absence of visible precipitate and stable optical density reading on nanodrop. Stocks were aliquoted using sterile glass syringes into argon purged, low binding tubes, sealed, and stored at –80°C. Aliquots were single use and were never subjected to repeated freeze thaw cycles. Under these handling conditions, anandamide retained biological potency in downstream assays.

### Animal Behavior

#### Delayed Non-Match to Place (DNMP)

The test was conducted in a T-maze with 3 arms (12in L x 3in W x 5in H) divided by sliding guillotine doors. In addition, the starting arm has an additional sliding door that defines a small compartment (3 x 3 x 3 in) used as starting box as well as “delay” box (see below). Mouse behavior was recorded using a ceiling-mounted camera under red light illumination and, for the photometry cohorts, analyzed using the Ethovision XT (Noldus) software.

A week prior to the start of the experiment all food was removed from the homecages and mice started a food deprivation regime to reach 80-85% of their initial weight. During this week, mice were daily handled and weighted, they were given ∼1.5g of regular chaw/mouse and habituated to chocolate pellets (BioServ cat F05301). Then, they were habituated to the T-maze over 2-3 d before DNMP training. During habituation, mice were allowed to freely explore the maze, where 20mg chocolate pellets were placed at the end of each arm, for 20-30 min. In the case of the photometry experiments, mice were habituated with the recording tether.

On the subsequent days, mice underwent DNMP training (for typically 3-4 days, until individual animals reached performance criteria of at least 75% accuracy. Group means tend to be 80-90%). Training consisted of 20 trials per day, where each trial has two phases, a forced choice followed by a free choice. During the forced choice, mice could only visit one arm, provided with one chocolate pellet at the end of it, while the access to the other arm was blocked by the door. The mouse was then enclosed in the delay box at the end of the starting arm for 5 sec before the door is lifted to start the free choice. In this phase, both arms are open, and only the one opposite to the arm previously visited during the forced phase has a chocolate pellet. If the mouse enters the rewarded arm and eats the chocolate pellet the trial terminates and the mouse is moved to a holding area for the ITI phase (∼20sec) while the experimenter sets up the maze for the successive trial. During the inter-trial interval (ITI), the maze was wiped with 10% ethanol to remove potential scent cues left by previous trials, and chocolate pellets were re-baited. A trial was considered correct if the mouse entered the goal arm not visited during the forced choice and was able to collect the pellet, and incorrect if they revisited the same arm, where they were confined without reward for 20 s before being placed in ITI. The order of rewarded arms is pseudo-randomly assigned on a trial-by-trial basis. After training, all mice that reached performance criterion were submitted to a test phase, which had the same structure as the training phase (forced choice + delay + free choice + ITI) with the difference of longer delays (either 15s, or 30s or 60s) alternating in an interleaved fashion over a total of 30 trials in order to have a final count of 10 trials for each delay period.

For the pharmacology experiments, mice were habituated to i.p. saline injections daily prior to the start of the training as well as during the training (before the start of every training session). During the test, mice were injected with either saline or anandamide (Enzo Life Sciences, cat. no. BML-FA017) (diluted in saline and injected i.p. at 5mg/kg at a 10ml/kg volume)2 min prior to the beginning of the test session, and every 10 trials (∼20 min) for a total of 3 injections. In these cohorts, mice were submitted to two consecutive test days where each day half of the group received saline and the other half received anandamide. The pharmacological treatment was inverted on the following day.

For photometry recordings, mice injected with camKIIa-GFLAMP2 and implanted with fiber optical cannulas, were recorded at different time points after surgery (day 10, day 25) to measure baseline GFLAMP2 signal and ∼40 days after surgery to record GFLAMP2 signals during behavior. For these experiments, the optical cannulas of mice were wiped with 10% ethanol and tethered to optical fibers (BBP_400/430/1100-0.57_4.5m_SMA-4xZF1.25_LAF, Doric Lenses), and left in the homecage for baseline recordings (10 minutes), or introduced in the maze to perform the behavioral experiment (habituation, training and test). The mouse behavior during the DNMPT photometry experiments was recorded by a ceiling mounted camera and analyzed by the Ethovision XT (Noldus) software to record both the motor activity and their position in the maze.

#### 5 arm Radial Maze

The radial arm maze consists of a central hub with 5 equidistantly spaced arms radiating outwards (17in L x 3in W x 6in H, black) on a white floor. Each arm, except for one, has blue tape on the floor arranged in a distinct pattern, to provide visual and tactile cues (see insert in Extended Data Fig. 7c). Mice were handled for 4 days and habituated to the i.p. saline injection. On the test day, mice were acclimatized to the experimental site for 1 h before the experiment started, then injected with either saline or anandamide (5mg/kg, i.p.) 2 minutes before being introduced in the maze one at the time and let free to explore it for 15 minutes. The mouse behavior was recorded with a ceiling-mounted camera and tracked with Ethovision XT (Noldus). The motor activity was measured to rule out arm biases and other locomotor changes between genotypes and drug groups that can confound task performance; this was assessed as distance moved (in inches) automatically calculated by the software, as well as the number of entries, defined as the mouse having all 4 paws inside the arm, manually scored by an experimentalist blind to the groups. The sequence of arms visited was used to measure correct alternations defined as the mouse visits three, four or five new arms before re-entering an arm already visited (e.g.: “ACDEB”). Short-term memory is assessed as percent of correct 3-, 4-, or 5-alternation sets over total alternations made.

### *In Vivo* Fiber Photometry

#### Photometry System

A custom multi-fiber photometry system was built as previously described in Hsiao et al., 2020. For G-FLAMP2 imaging, 470-nm excitation light was provided by an LED (Thorlabs M470F3) and routed through a dichroic mirror holder containing a 425-nm long-pass filter (Thorlabs DMLP425R), then reflected by a 495-nm long-pass dichroic mirror (Semrock FF495-Di02-25×36). Excitation light was focused through a 20×/0.5-NA objective (Nikon CFI S Fluor 20×, MRF00100) onto custom-length, branching, low-autofluorescence fiber-optic patch cords composed of 400-µm-diameter, NA 0.57 optical fibers (Doric Lenses BBP(4)_400/430/1100-0.57_4.5m_SMA-4×ZF1.25_LAF). Patch-cord termini were coupled to custom-length fiber cannula implants (Doric Lenses MFC_400/430-0.48_3.2mm_ZF1.25_FLT) using ceramic mating sleeves (SLEEVE_BR_1.25). Fluorescence emission from thalamic neurons was collected through the same fiber pathway and passed through a GFP emission filter (Semrock FF01-520/35-25) before amplification and imaging onto a high-sensitivity sCMOS camera (Prime 95B, Photometrics). An Arduino Uno microcontroller was used to trigger synchronized photometry acquisition and behavioral position tracking via EthoVision XT v14 software.

#### Photometry data processing and analysis

Raw photometry signals, captured at ∼20 Hz, were processed and analyzed using custom MATLAB code. Regions of interest were manually drawn in the center of each optical fiber to extract fluorescence. The raw fluorescent intensities extracted from the ROIs were processed similar to what has been previously described^70^.

Briefly, for each manually defined ROI, the raw fluorescence time series (F_unsplit_raw) was extracted across all frames. Signals were low-pass filtered using a zero-phase, sixth-order Butterworth filter (half-power frequency, 0.5 Hz; sampling rate, 12 Hz) implemented with forward–reverse filtering (filtfilt) to avoid phase distortions. To correct for slow baseline drift (e.g., photobleaching), the filtered trace was fit with a biexponential model (exp2), and the residuals from this fit were used as the baseline-corrected fluorescence (F_e_bsub). Baseline-corrected traces were then z-scored across the entire session (F_e_z). This z-scored traced was used for all downstream analyses. Metadata (animal identifier, recording date, region, frame count, ROI count) and fit parameters/goodness-of-fit were stored for each ROI alongside the processed signals.

Photometry signals were synchronized and aligned to task related events (cue, delay, choice, ITI) through time-stamps of behavioral frames captured through Ethovision. The change in cAMP during the delay period (ΔcAMP Delay) was computed by subtracting the mean signal from the first 250ms and last 250ms of the delay period. This approach captured the net change in G-Flamp2 fluorescence across the delay (memory maintenance) period. ΔcAMP delay values were calculated on a per-trial basis for each mouse, then averaged within each mouse to obtain per-animal means. A similar process was used to calculate the change in G-Flamp2 fluorescence during the ITI (ΔcAMP ITI). Significant differences in these measures between genotypes were assessed using one-way ANOVA.

### *In Vivo* Calcium Imaging

#### Surgery & Behavior

Mice were injected with 800 μL of AAV1-CaMKIIa-GCaMP6f (Addgene 100834; 1.0×10^13^ vg/mL) into PFC (AP: +2.0, ML: –0.35, DV: –1.7). Immediately following this injection, a GRIN lens of 0.6 mm diameter, 7.3mm length, and 0.5 NA (Inscopix 1050-004597) was implanted above the injection site (AP: +2.0, ML: –0.35, DV: –1.5) and sealed with optic glue. Finally, a guide cannula (RWD 62003) was implanted above the mediodorsal thalamus (AP: –1.4, ML: –0.4, DV: –2.9). A custom titanium headplate was glued to the skull, and all implants were fixed in place with dental cement. After recovery from surgery, mice began DNMP training (5s delay periods). On test days (30s delay periods), mice were injected with 250 μL of sterile PBS or 288 μM AEA at 100 μL/min through an internal cannula (RWD 62203) extending 0.5mm beyond the guide cannula, and behavior commenced immediately.

#### Imaging & Analysis

In vivo calcium imaging was performed as previously described in Toader et al., 2023. Briefly, the mice were headfixed and each GRIN lens was cleaned with 100% methanol. Custom 3D printed adapters were used to couple each lens to a bundle of imaging fibers with diameter 0.6mm and core-to-core distance of 3.3 μm (Myriad Fiber, FIGH-30-650s). Imaging was achieved by excitation through a 473 nm laser (0.25 mW laser power at the end of the fiber tip) and imaging using a CMOS camera (Photometrics Prime 95B). Images were collected using the Photometrics data acquisition software, Programmable Virtual Camera Access Method (PVCAM), at 34 Hz and 920 pixel/mm.

Source extraction was performed as previously described in Toader et al., 2023. The field of view was cropped and motion-corrected using NoRMCorre, Pnevmatikakis and Giovannucci, 2017. A combination of CNMF-E, Zhou et al., 2018, and custom packages were used to identify and isolate individual cell ROIs and dF/F signals; each extracted cell was manually validated. Cell registration across sessions was performed with CellReg, Sheintuch et al., 2017. For each detected cell, its normalized ΔF/F was calculated by dividing (F-F_baseline_)/F_baseline_, where F is the raw fluorescence and F_baseline_ is the mean of the fluorescence value for that cell over a 20s sliding window in order to account for changes across the recording session. To identify significant transients, we identified a noise estimate as previously described in Rajasethupathy et al., 2015 and defined transients as events exceeding 3σ of the estimated noise level for at least 300 ms (approximately twice the duration of the half-life decay time of GCaMP6f).

Epoch tuning was assessed for each neuron by a shuffle-based permutation test: each neuron’s full-session transient vector was randomly circularly permuted 1000 times, preserving total event count and relative event timing while destroying behavioral timing. Observed epoch event counts were then compared with the neuron-specific shuffled null distribution. Neurons were classified as epoch excited if their observed count exceeded the 95th percentile of the null distribution.

For visualization of trial averaged GCaMP traces, because the timing of cue and free choice events differed across trials, a standardized pseudotrial structure was used, in which we sampled 20s of inter-trial interval (ITI), 2s of the cue (forced choice), the 30s memory period, and 2s of the free choice period.

#### URB-597 effects on working memory performance in macaques

Rhesus monkeys (n=14, 12-30 years, 2 male, 12 female) were pretrained on the variable delay spatial delayed response task, a test of visuospatial working memory performed with a manual response in a Wisconsin General Test Apparatus. Monkeys test for highly palatable food rewards, limiting the need for food regulation. Each daily session consisted of 30 trials with 5 different delays ranging from 0 sec to the delay in which that monkey performed at chance, thus allowing the opportunity for drug to either improve or impair performance. Stable baseline performance of about 70% correct was established for all animals. Monkeys were tested by an experimenter who was highly familiar with the normative behavior of each animal, but unaware of drug treatment conditions. Animals were rated for changes in sedation/agitation and aggression using 9 point rating scales. The FAAH inhibitor, URB-597, or vehicle control, was administered intramuscularly 2 hrs before testing. Pilot doses between 0.0001-0.1 mg/kg were initially explored; an optimal dose range between 0.001-0.03 mg/kg was the focus of subsequent testing compared to vehicle control. There were washout periods of at least 10 days between doses; monkeys were required to return to stable baseline performance on vehicle prior to additional drug treatments. Statistical analyses employed a repeated-measures t-test with two-tailed significance. This study was carried out in accordance with the Guide for the Care and Use of Laboratory Animals issued by the National Institutes of Health (NIH), USA.

**Supplementary Figure S1.**
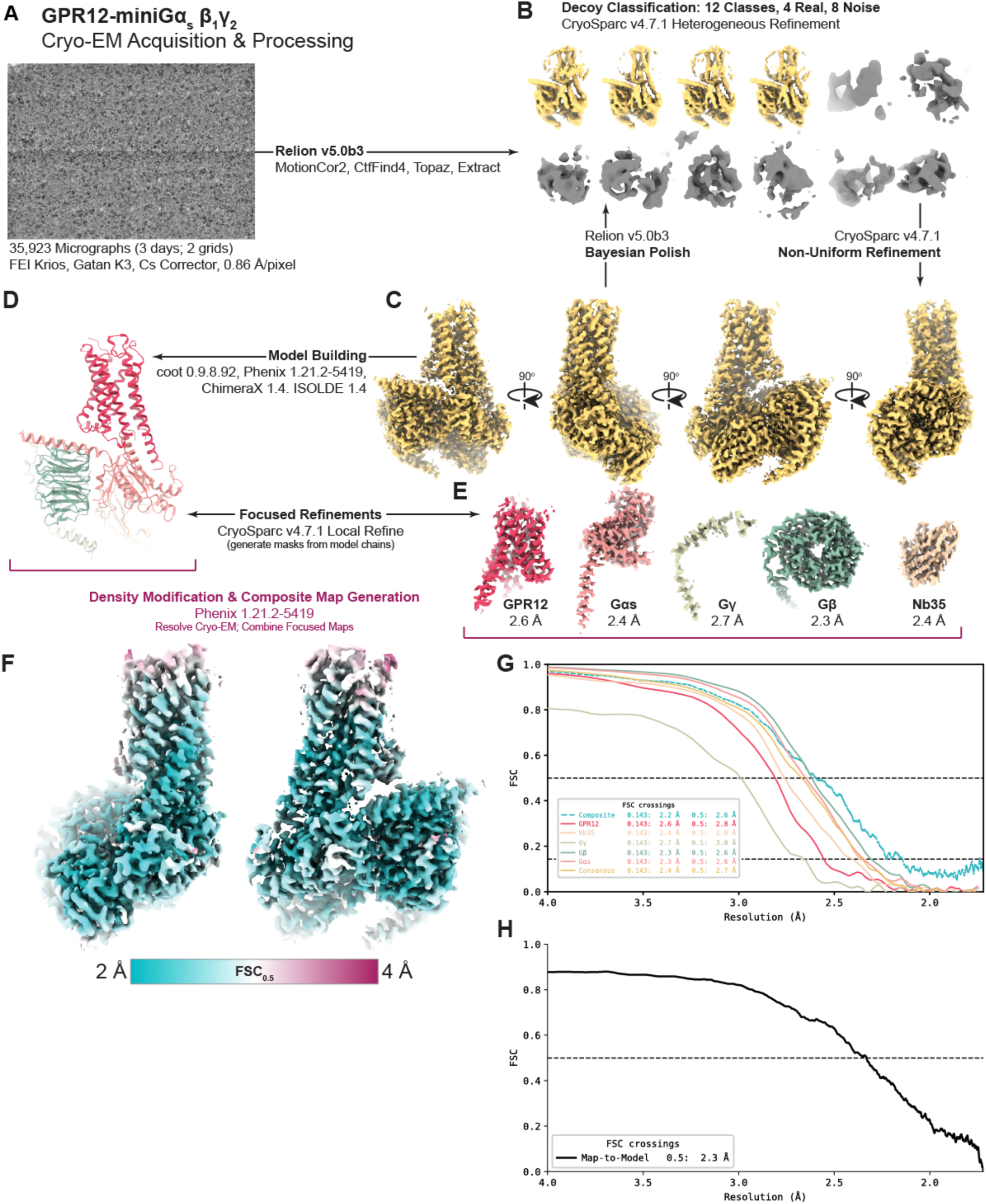
Generation of a cryo-EM structure of hGPR12. (**A**) Representative micrograph collected on a Titan Krios cryo-electron microscope. (**B**) Two-dimensional particle classifications, including structural and decoy classes, used for downstream refinement. (**C**) Cryo-EM density map after iterative 3D refinement. (**D**) Initial model building guided by the refined density. (**E**) Focused refinement of the heterotrimeric signaling complex and Nb35. (**F**) Local resolution map of the final reconstruction. (**G)-(H),** Fourier shell correlation (FSC) curves showing resolution determination for individual subunits (**G**) and the map-to-model comparison (**H**).

**Supplementary Figure S2.**
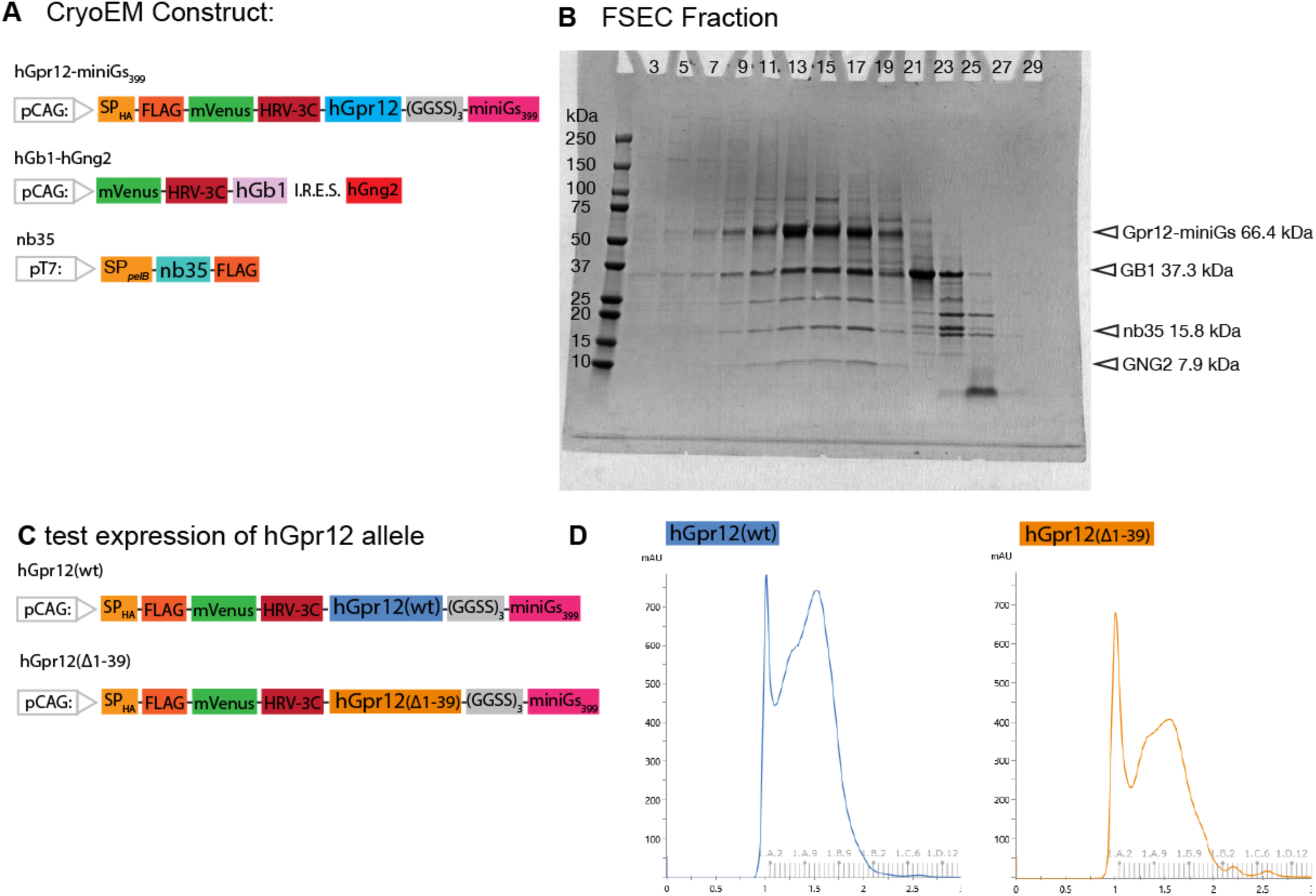
Construct design, expression screening, and purification of hGpr12 for cryo-EM studies. (**A**) Schematic representation of the cryo-EM expression constructs used for structure determination. The Gpr12–miniGs signaling complex was assembled using N-terminally FLAG-tagged human Gpr12 fused to miniGs, together with Gβ1 and Gγ2 subunits. Additional expression constructs for Gβ1 and Gγ2 are shown. (**B**) Coomassie-stained SDS-PAGE analysis of size-exclusion chromatography fractions from purification of the Gpr12–miniGs signaling complex. Fractions corresponding to the purified complex used for cryo-EM studies are shown. Bands corresponding to Gpr12–miniGs (66.4 kDa), Gβ1 (37.3 kDa), and Gγ2 (7.9 kDa) are indicated. (**C**) Schematic representation of Gpr12 constructs used for expression screening, including wild-type Gpr12 and an N-terminal truncation mutant (Δ1–39) fused to miniGs. (**D**) Representative fluorescence-detection size-exclusion chromatography (FSEC) profiles of wild-type Gpr12 (left) and the Gpr12 Δ1–39 truncation construct (right). Both constructs produced a monodisperse receptor population; however, the Δ1–39 construct exhibited reduced expression yield relative to wild-type Gpr12. Consequently, the full-length receptor was selected for subsequent purification and cryo-EM studies.

**Supplementary Figure S3.**
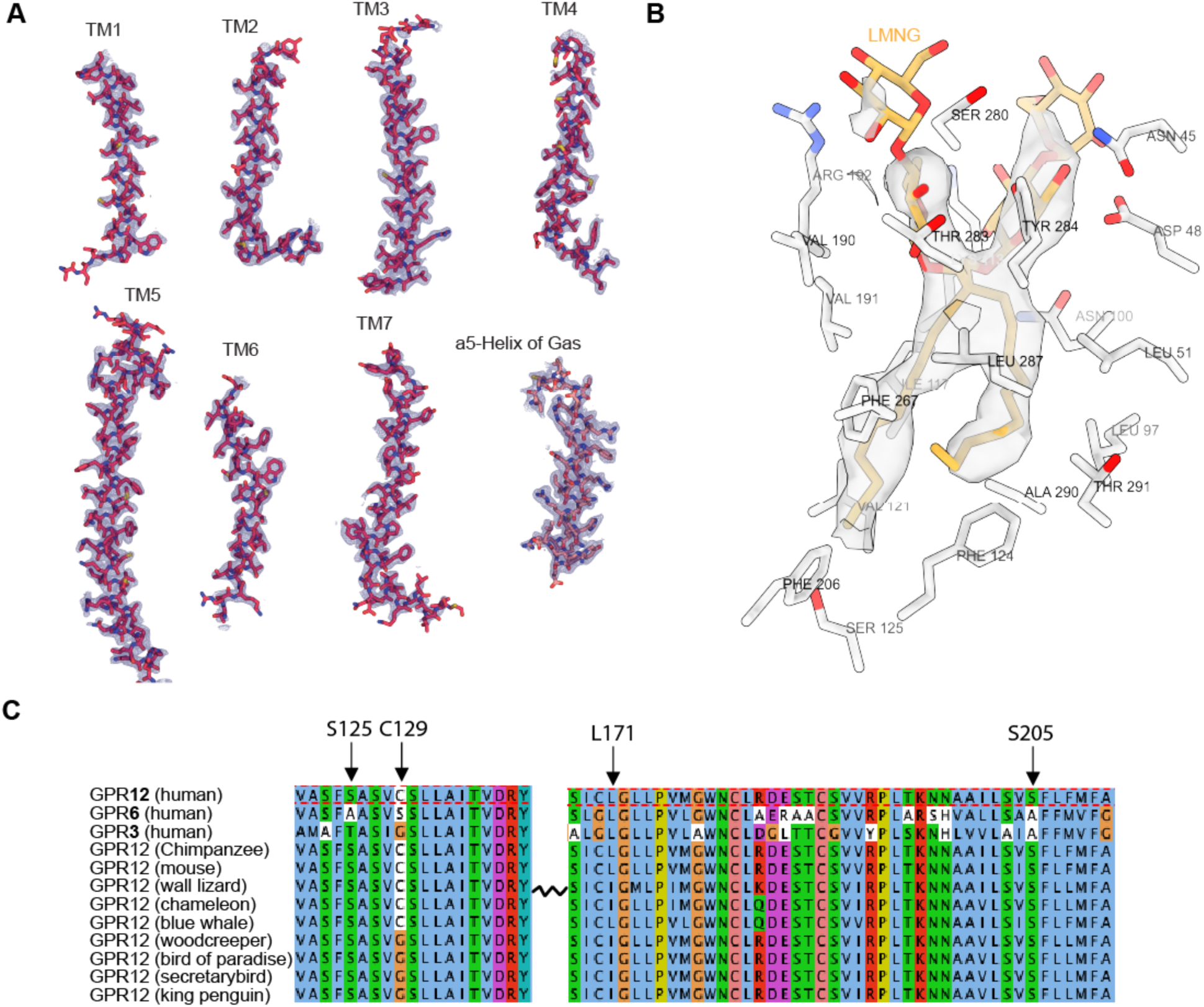
Computational, and functional characterization of Gpr12. (**A**) Overlay of the cryo-EM density map (purple) with the primary amino-acid sequence of Gpr12 (red). (**B**) Cross-sectional view of the orthosteric pocket exhibiting coordination of LMNG (yellow) by orthosteric residues (white) with cryo-EM density overlaid (light grey). (**C**) Alignment of hGPR12 (top), hGPR6 (second row), hGPR3 (third row), and cross-species Gpr12 alleles (Below). KS5 pocket residues S125, C129, L171, and S205 are indicated. Coloring is based on physiochemical amino acid characteristics. Uneven line in-between alignment blocks signify abridging primary amino acid sequence for visualization.

**Supplementary Figure S4.**
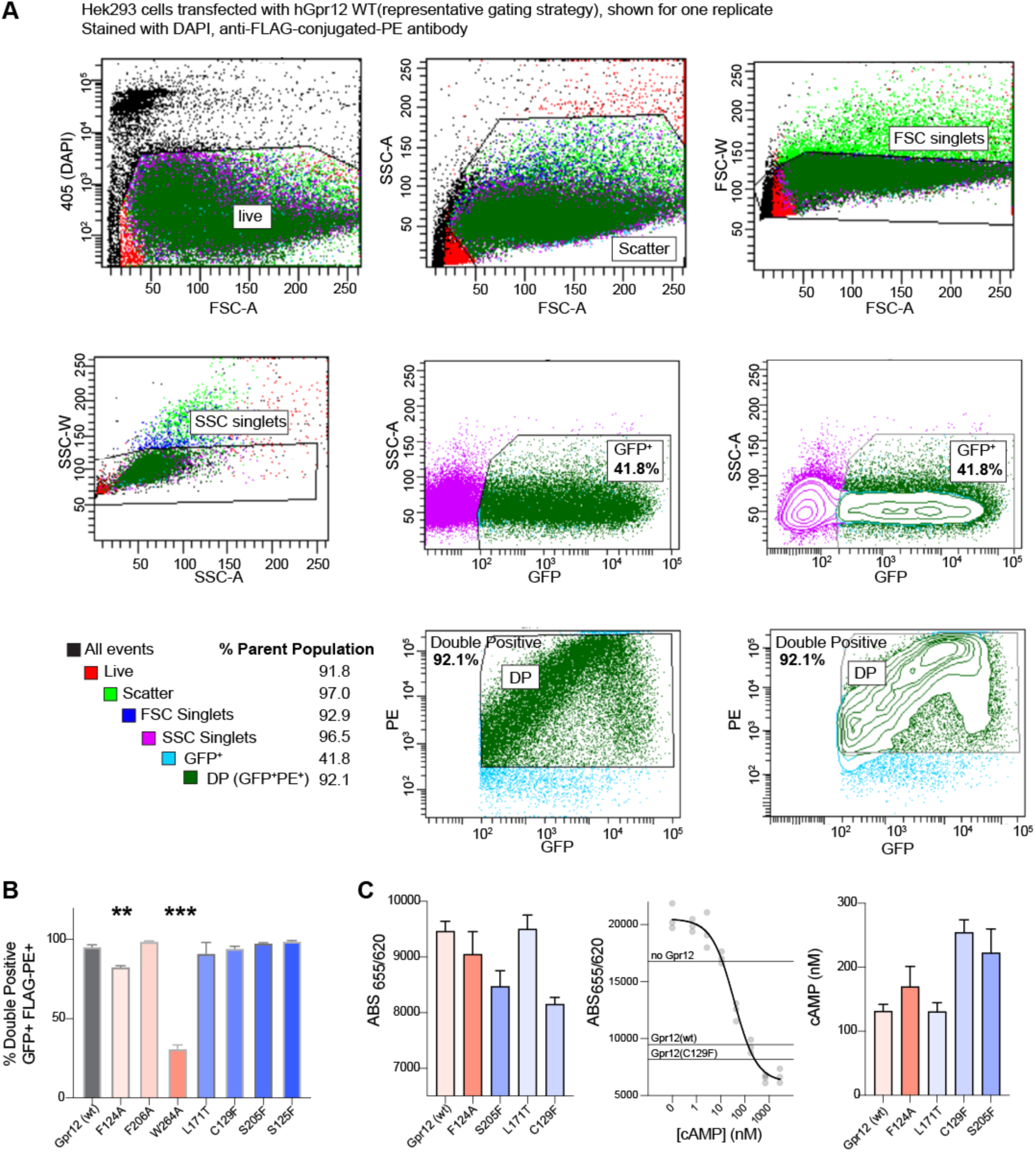
Cell Surface expression of hGpr12 Alleles. (**A**) Representative FACS gating strategy to measure percentage of cell surface expression. Number of PE+GFP+ cells (double positive, 92.1%) are calculated out of all GFP+ cells (41.8% of live cells (SSC singlets)). Percent of parent populations are depicted on the left. (**B**) Raw percentage of double positive GFP+ PE+ cells over total GFP+ cells per genotype. Data are mean +/− SEM (n=3 biological replicates), ** p< 0.01, *** p<0.001 one-way ANOVA with Dunnett’s test. (**C**) cAMP accumulation assay data processing used in Figure 1h. The FRET signal ratio between d2-labeled cAMP (665-nm emission) and a Europium Cryptate-labeled monoclonal cAMP antibody (620-nm emission) was measured (left). This ratio exhibits a linear relationship with cAMP concentration (center), enabling interpolation of absolute cAMP values using a standard calibration curve (right). Data are mean n=3 biological replicates.

**Supplementary Figure S5.**
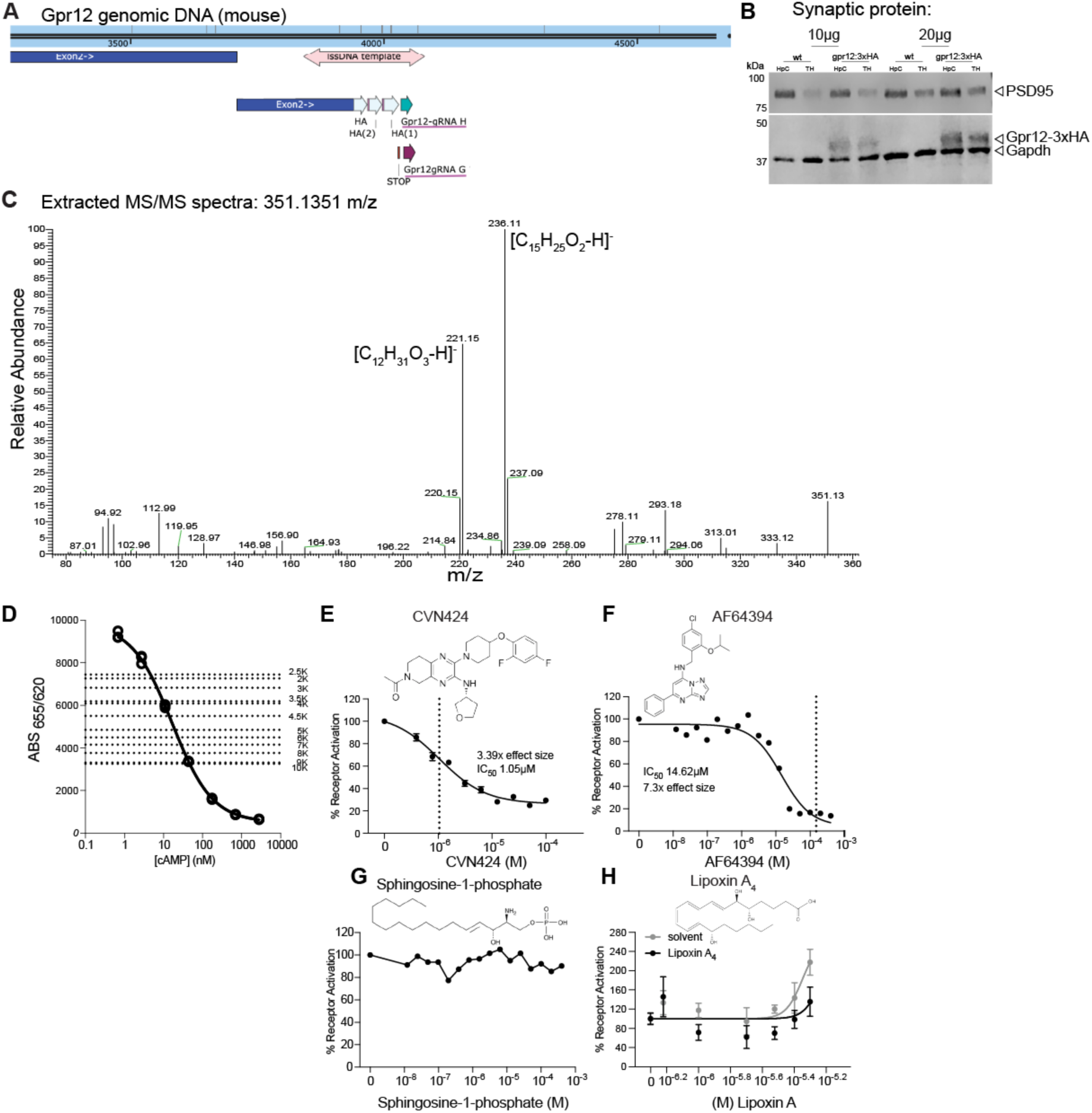
Generation of the Gpr12:3xHA mouse line, lipidomics assignment, and validation of the doxycycline-inducible assay system. (A) Genomic alignment of mouse *Gpr12* exon 2 illustrating the CRISPR knock-in strategy for endogenous 3×HA tagging. A single-stranded DNA donor template (salmon) inserts a tandem 3xHA epitope at the C terminus of *Gpr12* exon 2 (dark blue) following a double stranded break generated by two guide RNAs, Gpr12-gRNA H (green) and Gpr12-gRNA G (purple). (B) Western blot validation of Gpr12:3×HA expression in 10ug or 20 ug synaptic fractionations of Hippocampus (HpC) and Thalamus (TH) tissue compared to C57 (wt) control mice pooled from 5 individual mice. HA, Psd95 and Gapdh bands shown. Expected molecular weights are 40kDA for Gpr12:3xHA, 95kDa for Psd95, and 36kDA for Gapdh. (C) MS/MS spectra of fragmentation of the 351 m/z species identified in Fig. 2b, consistent with a 20:4 eicosanoid backbone. Fragment ions 221.15 m/z and 236.11 m/z are assigned [C12H_31_O3-H]^-^ and [C_15_H_25_O_2_-H]^-^, respectively. (D) Validation of the HTRF cAMP assay using serial dilutions (between 2.5K and 10K) of HEK293T cells expressing hGpr12 (dashed lines) plotted against the manufacturer-defined linear dynamic range of the assay (bold lines). (E)-(F), Pharmacological validation of the assay using known Gpr12 inhibitors CVN424(n=2) (E) and AF64394(n=1) (F). (G)-(H), Additional ligand testing using Sphingosine-1-phosphate(n=1) (G), and the eicosanoid Lipoxin A4 (n=4) (H). Data are expressed as percent activation over the solvent, except for H where we observed a solvent effect at the higher concentrations. All data are shown as mean +/− SEM.”

**Supplemental Figure S6.**
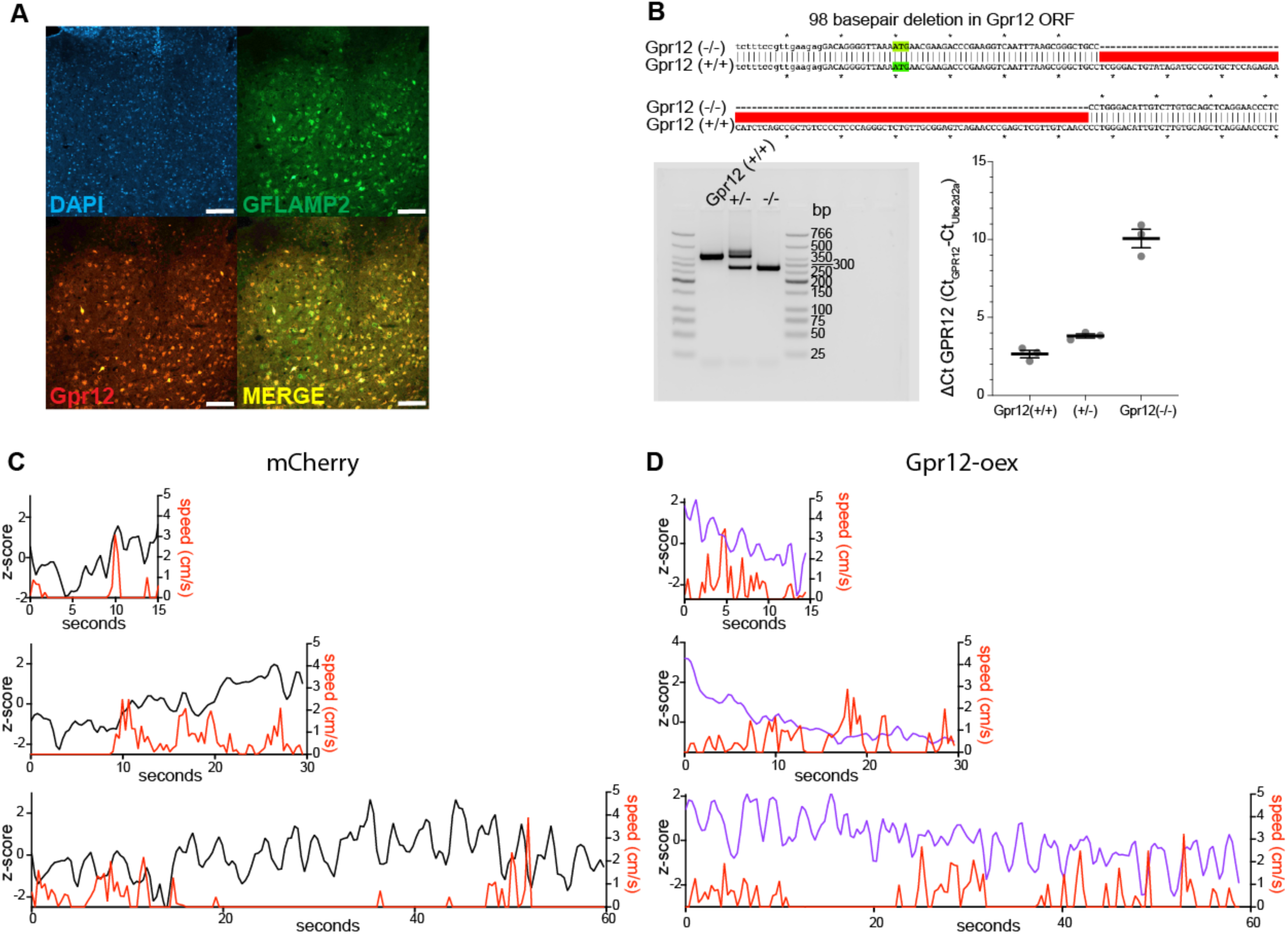
Histological validation, Gpr12 knockout generation, and behavioral analyses related to figure 3. (**A**) Histological validation of photometry cohort mice from Fig. 3c showing mCherry co-expression with the GFLAMP2 cAMP sensor (green) and Gpr12 (red). Scale bar 100 µm (inset) 20x. (**B**) Sequencing validation of the 98-bp genomic deletion generating the Gpr12 knockout (top), with PCR confirmation (bottom left) indicating loss of the expected DNA amplicon in Gpr12(−/−) animals, and qPCR showing Gpr12 mRNA expression in the mediodorsal thalamus of wildtype (Gpr12(+/+)), het (Gpr12(+/−)) and knockout (Gpr12(−/−)) mice, n=3/each with Gpr12 mRNA ct values normalized to the abundant Ube2d2a mRNA. Data are mean +/− SEM. (**C), (D**) Representative single trials from C) an mCherry control and D) a Gpr12-overexpressing (Gpr12-oex) animal showing 15 (top), 30 (middle), and 60s (bottom) delay periods, with running speed (red) and photometry signal (mCherry, black and Gpr12-oex, purple).

**Supplemental Figure S7.**
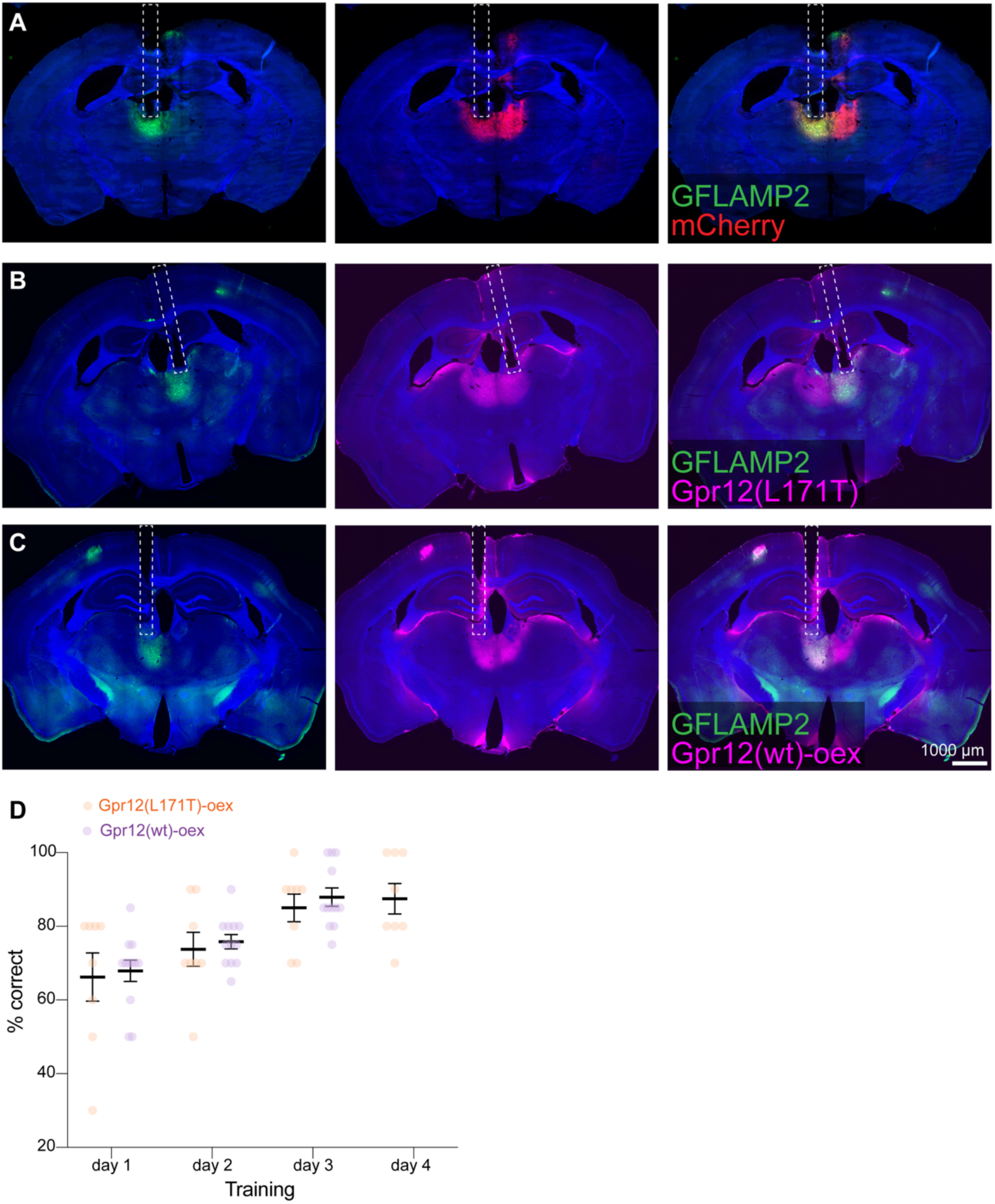
DNMP training curve and histological validation of viral expression and cannula placement related to photometry Figures 4. (**A**) Coronal sections of viral expression in the mediodorsal thalamus showing GFLAMP2 expression (left), mCherry expression (center), and GFLAMP2/mCHerry overlay (right) with unilateral cannula placement (white dash). (**B**) Coronal section of animal expressing GPR12(L171T) constrct (purple) with same organization and scale as (**A**). (**C**) Coronal section of animals expressing GFLAMP2 (left), Gpr12(wt)-oex (purple, center), with overlay and cannula track placement (right). Scale bars: 1000 µm (coronal) 4x stich, cannula diameter, 400 µm.(**D**) Learning curve during the training phase of the DNMP task in Gpr12(L171T)-oex (n=8, orange) and Gpr12(wt)-oex (n=12, purple). Data are mean +/− SEM. Two-way ANOVA with repeated measures, n.s. The Gpr12(L171T)-oex group received an extra day of training in the attempt to bring a few borderline mice to criteria. Subsequently, the mice will be submitted to the test shown in Fig. 4H.

**Supplemental Figure S8.**
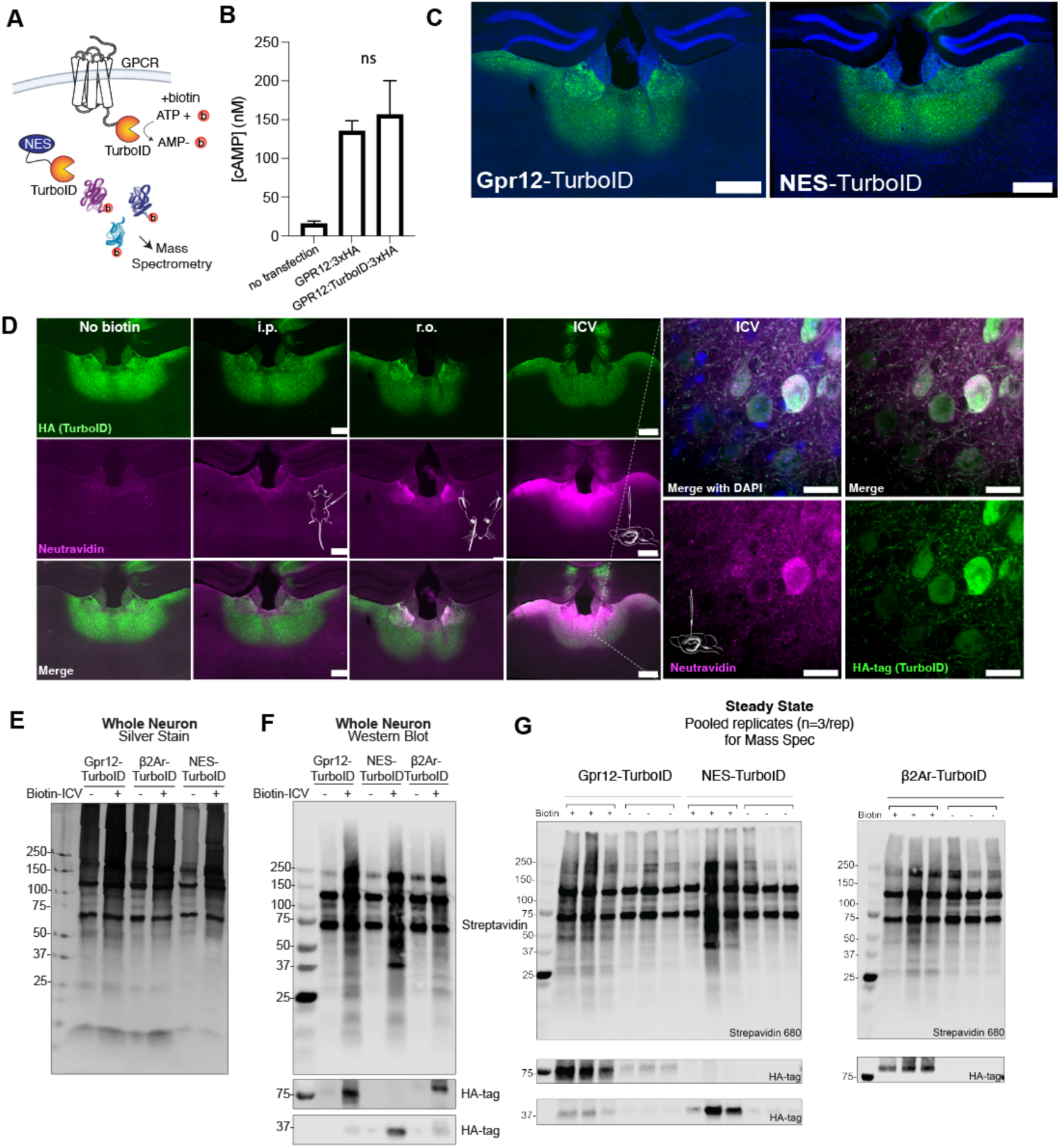
Validation of Gpr12-TurboID labeling and proteomic workflow. (**A**)TurboID approach, a C-Terminal fusion of TurboID labels proximal proteins with biotin.(**B**) In vitro cAMP accumulation assay comparing Gpr12-3xHA and Gpr12-TurboID-3xHA expressed in HEK293 cells. Fusion of TurboID to Gpr12 did not significantly alter receptor signaling. Data are mean ± SEM from n = 3 independent experiments. (**C**) Coronal sections of viral expression in the mediodorsal thalamus showing expression of Gpr12-TurboID and control NES-TurboID (green) Scale bars, 100 μm. (**D**) Evaluation of biotin delivery routes for in vivo TurboID labeling. Representative images of mediodorsal thalamus expressing Gpr12-TurboID following no biotin treatment, intraperitoneal (i.p.), retro-orbital (r.o.), or intracerebroventricular (ICV) biotin administration. HA immunoreactivity (green) marks Gpr12-TurboID expression and NeutrAvidin labeling (magenta) indicates biotinylated proteins. Higher-magnification images from the ICV condition demonstrate robust local biotinylation surrounding Gpr12-TurboID-expressing neurons. Scale bars, 100 μm (left panels) and 10 μm (right panels). (**E**) Silver-stained gel of whole-neuron lysates from Gpr12-TurboID, β2AR-TurboID, and NES-TurboID control groups with and without ICV biotin administration. Samples were used to assess overall protein recovery and labeling conditions prior to downstream proteomic analysis. (**F**) Western blot analysis of whole-neuron lysates probed with streptavidin and HA antibodies. Biotin administration produced robust protein biotinylation in TurboID-expressing samples while confirming expression of the corresponding HA-tagged constructs. (**G**) Streptavidin and HA immunoblots from pooled biological replicates used for mass spectrometry. Three independently pooled samples (n = 3 pools per condition) were generated for Gpr12-TurboID, β2AR-TurboID, and NES-TurboID groups under biotin-treated and untreated conditions. Comparable construct expression and biotin-dependent labeling were observed across samples submitted for proteomic analysis.

**Supplementary Figure S9.**
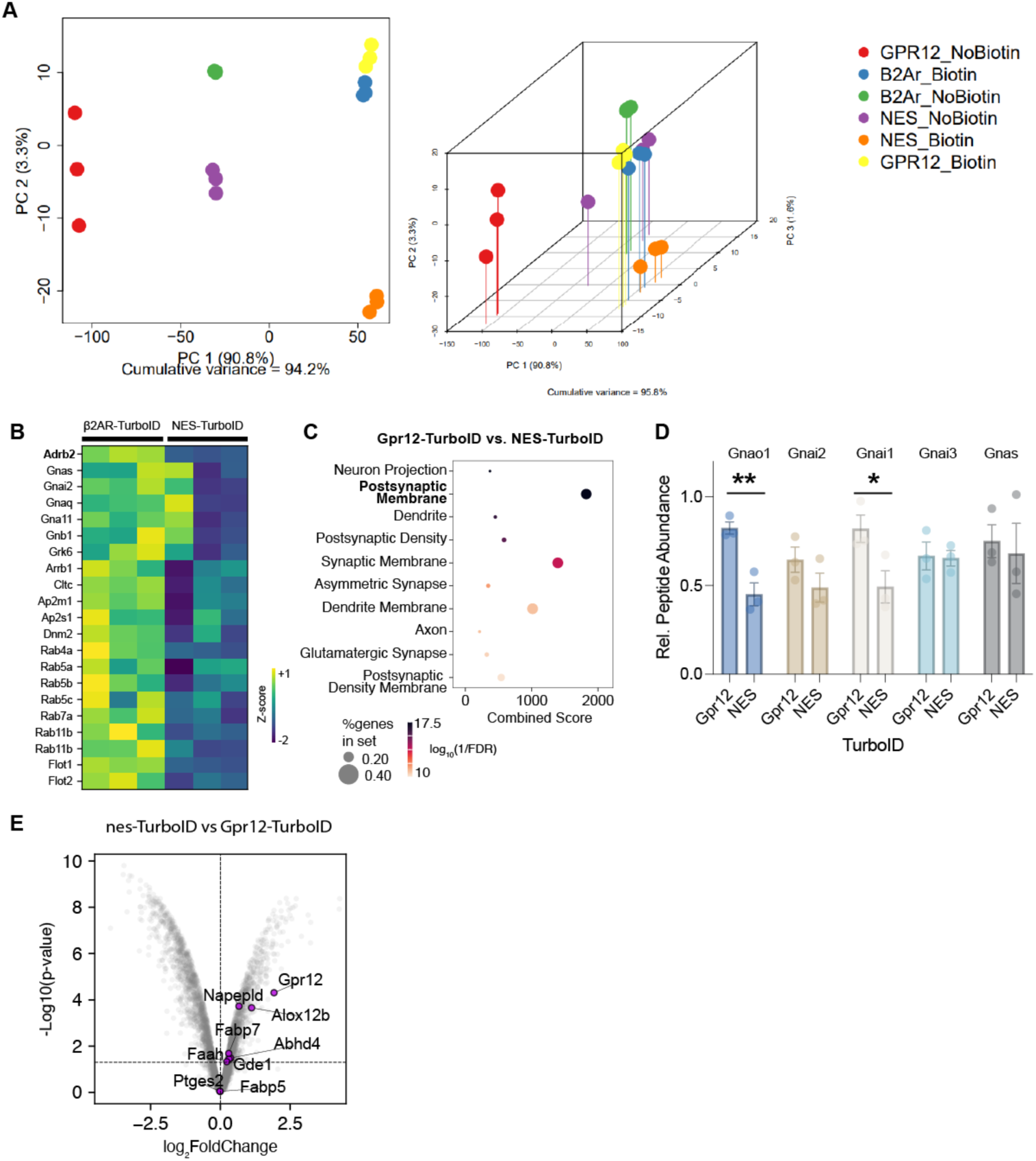
Proteomic quality control and validation of Gpr12-TurboID proximity labeling. (**A**) Principal component analysis (PCA) of TurboID proteomic datasets. Two-dimensional (left) and three-dimensional (right) PCA plots demonstrate separation of Gpr12-TurboID, β2AR-TurboID, and NES-TurboID samples in the presence or absence of biotin. Biological replicates cluster tightly within groups, indicating high reproducibility of proximity-labeling and mass spectrometry measurements. (**B**) Heatmap showing relative abundance of proteins previously identified in β2AR proximity-labeling datasets. Protein abundances are shown as z-scored values across β2AR-TurboID and NES-TurboID biological replicates, demonstrating enrichment of established β2AR-associated proteins in β2AR-TurboID samples. (**C**) Gene ontology (GO) enrichment analysis of proteins identified in Gpr12-TurboID relative to NES-TurboID controls. The top enriched cellular component terms are associated with postsynaptic, dendritic, and synaptic membrane compartments, consistent with localization of Gpr12 within excitatory neuronal processes. (**D**) Relative peptide abundance (TMT ratios) of the G protein α-subunits Gnao1, Gnai1, Gnai2, Gnai3, and Gnas in GPR12-TurboID versus Nes-TurboID control proteomic datasets. Data are shown as mean ± SEM. Statistical significance is reported as FDR-adjusted p values derived from the full proteomic comparison presented in (E). (**E**) Volcano plot comparing proteins identified in Gpr12-TurboID and NES-TurboID samples. Highlighted proteins include Gpr12 and multiple enzymes involved in endocannabinoid metabolism and lipid signaling, including Napepld, Abhd4, Gde1, Faah, Faah2, Fabp5, Fabp7, and Alox12b. Differential enrichment of these proteins supports the association of Gpr12 with endocannabinoid biosynthetic and metabolic pathways within the mediodorsal thalamus.

**Supplementary Figure S10.**
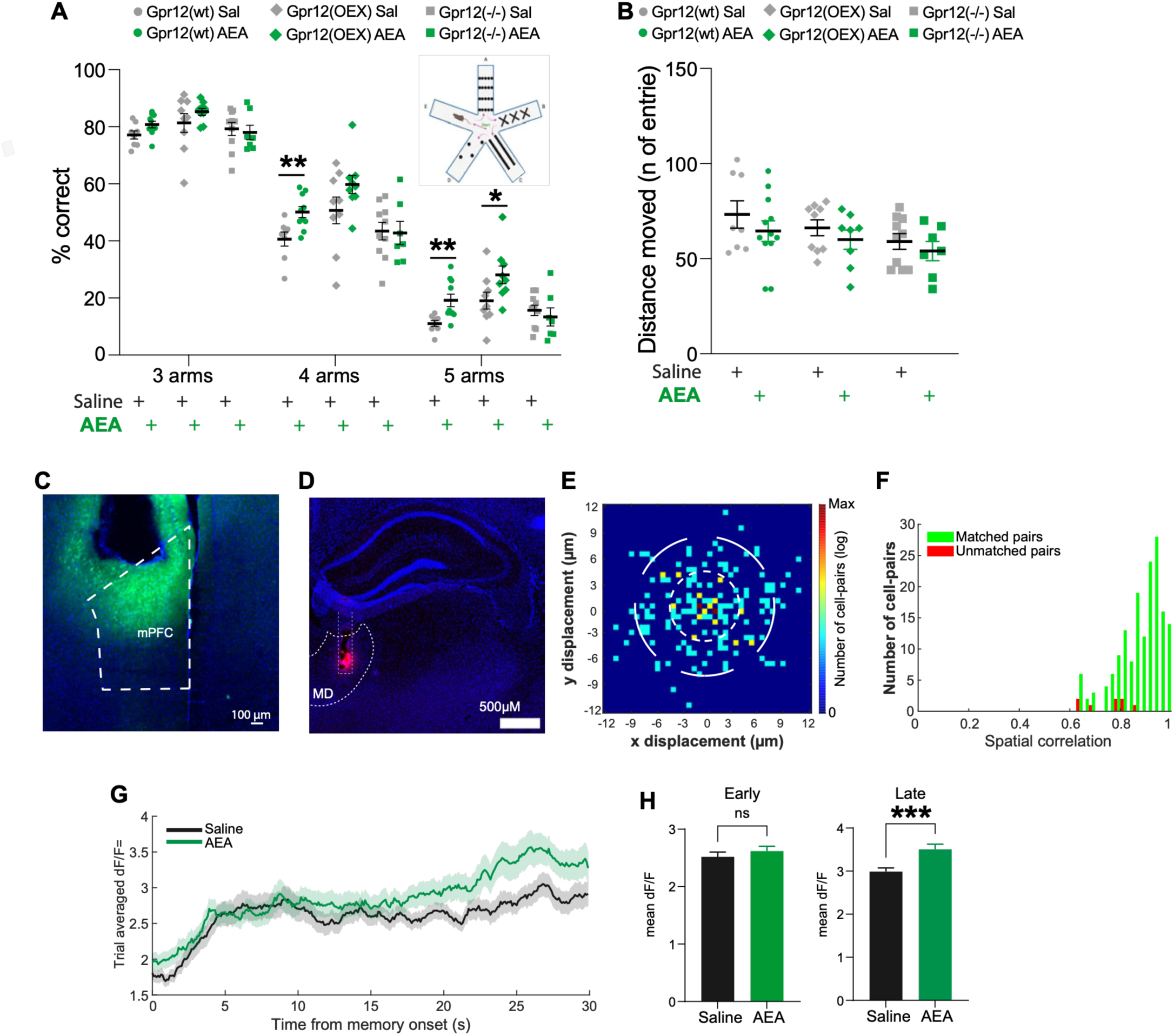
Effects of anandamide on behavior and prefrontal neural activity. (**A**) Performance in the five-arm spontaneous alternation task expressed as % correct 3-, 4-, and 5-arm alternations over the total alternations made during a 15-min task, in Gpr12 wt (circle), Gpr12-oex (diamond), Gpr12 knockout (Gpr12(−/−), square) injected i.p. with either Saline or 5mg/kg anandamide (AEA). Data are mean +/− SEM (n=8-10/each group), Welch’s t test * p < 0.05, ** p < 0.01. (**B**) Motor activity assessed during the five-arm spontaneous alternation task measured as total number of arm entries during a 15-min task. Welch’s t test n.s. (**C**) Representative coronal section of imaging site in PFC showing GCaMP6f expression and lens track. Scale bar=100μm. (**D**) Representative coronal section of injection site in MD showing restricted Dil injection. Scale bar=100μm. (**E**) Post-alignment spatial footprint offset map of registered PFC cell pairs across imaging sessions. (**F**) Distribution of spatial correlations for nearest-neighbor cell pairs showing matched (green) and non-matched (red) pairs. (**G**) Trial-averaged memory-period dF/F traces for all memory-activated registered neurons under saline and AEA conditions (n=86 neurons). Traces aligned to memory onset; shading indicates SEM. (**H**) Quantification of mean dF/F for all memory-activated neurons during early and late memory periods. AEA did not significantly alter early memory but increased late memory period activity (n=134 saline and 128 AEA neurons; unpaired two-tailed t test) ***=P<0.001.

#### Cryo-EM data collection, refinement and validation statistics

**Supplementary Table S1.** Cryo-EM data collection, refinement, and model validation statistics for the Gpr12– miniGs signaling complex. Summary of cryo-EM data acquisition parameters, image processing statistics, model refinement metrics, and structural validation statistics for the Gpr12–miniGs complex. Reported values include microscope and detector settings, particle numbers, map resolution, model composition, geometric quality metrics, and validation parameters used to assess the final cryo-EM reconstruction and atomic model.

|  |  |
| --- | --- |
|  | GPR12:Gs<br>+LMNG<br>(EMDB-xxxx)<br>(PDB xxxx) |
| **Data collection and processing** |  |
| Magnification | 105,000x |
| Voltage (kV) | 300 |
| Electron exposure (e-/Å<sup>2</sup>) | 50 |
| Defocus range (μm) | -0.8 to -1.6 |
| Pixel size (Å) | 0.86 |
| Symmetry imposed | C1 |
| Initial particle images (no.) | 3,237,839 |
| Final particle images (no.) | 1,267,653 |
| Map resolution (Å) | 2.4 Å (0.143) |
| FSC threshold |  |
| Map resolution range (Å) | 2.3 Å to 2.7 Å |
| **Refinement** |  |
| Initial model used (PDB code) | 7Y3G |
| Model resolution (Å) | 2.4 Å (0.5) |
| FSC threshold |  |
| Model resolution range (Å) | 2.3 Å to 2.7 Å |
| Map sharpening *B* factor (Å<sup>2</sup>) | 90.9 |
| Model composition | 16,445 atoms |
| Non-hydrogen atoms | 8,215 atoms |
| Protein residues | 1,031 |
| Ligands | 1 |
| *B* factors (Å<sup>2</sup>) |  |
| Protein | 58.15 |
| Ligand | 105.58 |
| R.m.s. deviations |  |
| Bond lengths (Å) | 0.005 |
| Bond angles (°) | 0.613 |
| Validation |  |
| MolProbity score | 0.85 |
| Clashscore | 1.23 |
| Poor rotamers (%) | 1.02 |
| Ramachandran plot |  |
| Favored (%) | 98.62 |
| Allowed (%) | 1.38 |
| Disallowed (%) | 0 |

